# Yeast with elevated chromosome numbers are addicted to high levels of Mps1

**DOI:** 10.1101/2023.01.09.523325

**Authors:** Régis E Meyer, Ashlea Sartin, Madeline Gish, Jillian Harsha, Emily Wilkie, Dawson Haworth, Rebecca LaVictoire, Isabel Alberola, Olivia Bowles, Hoa H Chuong, Gary J Gorbsky, Dean S Dawson

## Abstract

Tumor cell lines with elevated chromosome numbers frequently exhibit elevated expression of Mps1. These tumors are also dependent on high Mps1 activity for their survival. Mps1 is a conserved kinase involved in controlling aspects of chromosome segregation in mitosis and meiosis. The mechanistic explanation for the Mps1-addiction of aneuploid cells is unknown. To address this question, we explored Mps1-dependence in yeast cells with increased sets of chromosomes. These experiments revealed that in yeast, increasing ploidy leads to delays and failures in orienting chromosomes on the mitotic spindle. Yeast cells with elevated numbers of chromosomes proved vulnerable to reductions of Mps1 activity. Cells with reduced Mps1 activity exhibit an extended prometaphase with longer spindles and delays in orienting the chromosomes. One known role of Mps1 is in recruiting Bub1 to the kinetochore in meiosis. We found that the Mps1-addiction of polyploid yeast cells is due in part to its role in Bub1 recruitment. Together, the experiments presented here demonstrate that increased ploidy renders cells more dependent on Mps1 for orienting chromosomes on the spindle. The phenomenon described here may be relevant in understanding why high-ploidy cancer cells exhibit elevated reliance on Mps1 expression for successful chromosome segregation.

**AUTHOR SUMMARY:** Losing or gaining chromosomes during cell division leads to aneuploidy (an abnormal number of chromosomes) and can contribute to cancer and other diseases. Indeed, most cells in solid tumors carry abnormally elevated numbers of chromosomes. Mps1 is a regulator of the machinery that distributes chromosomes to daughter cells. In tumors with elevated chromosome numbers, the expression of Mps1 is often also elevated. In some aneuploid tumor cell lines these elevated Mps1 levels have been shown to be critical for tumor survival. To determine how cells with higher ploidy become dependent on Mps1, we explored Mps1-dependence in yeast cells with increased numbers of chromosomes. We report that yeast cells with elevated chromosome number are sensitive to reductions Mps1 expression. In cells with high ploidy and reduced levels of Mps1, the progression of the cell cycle is delayed and the ability of the cells to properly orient and segregate their chromosomes on the spindle is greatly reduced.

## INTRODUCTION

Maintaining error-free chromosome segregation during the cell cycle is critical for normal cell viability. Errors in chromosome segregation can lead to aneuploidy – an incorrect chromosome number - which is a common feature of cancer cells ^1^. To segregate correctly at anaphase, the sister chromatids of replicated chromosomes must attach to spindle fibers (microtubules) that emanate from opposite poles of the spindle ^2–4^. Microtubules connect to chromosomes at a proteinaceous structure called the kinetochore ^5^. The microtubule-kinetochore attachment process occurs in prometaphase. There is a stochastic aspect to the attachment process and in both yeast and mammalian systems many of the initial kinetochore-microtubule attachments are incorrect ^6–8^. Incorrect attachments include having the kinetochore of one chromatid attached to both spindle poles (something that doesn’t happen in budding yeast because there is only one microtubule per kinetochore) or having both sister chromatids attach to microtubules emanating from the same pole. These incorrect attachments are corrected by cycles of release and re-attachment of microtubules to the kinetochores. This process continues until each sister chromatid pair is properly attached to the spindle in a way that allows the sisters to segregate from one another at anaphase. Cells possess a surveillance system, called the spindle checkpoint, that delays progression into anaphase until all the sister chromatid pairs are properly attached to the spindle ^9^.

The conserved kinase Mps1 (also known as TTK in human cells) is one of the components of the chromosome segregation apparatus ^10–12^. This kinase is involved in multiple pathways during the cell cycle. The *MPS1* gene was initially identified for its essential role in duplication of the yeast microtubule-organizing center, known as the spindle pole body (SPB) (*MPS1* stands for monopolar spindle) ^13^. Subsequently Mps1 was found to act in both yeast and mammalian as a component of the spindle checkpoint where it phosphorylates Mad1 and, in budding and fission yeasts has also been shown to recruit Bub1/Bub3 to the kinetochores^14–18^. Mps1 also promotes re-orientation of chromosomes on the spindle and proper kinetochore microtubule attachment (in yeast and mammalian cells) both through its recruitment of Bub1, phosphorylation of other kinetochore components (e.g. the SKA, DAM1, NDC80 complexes, and Aurora B kinase), and probably other unknown mechanisms^19–23^ (reviewed in ^10^).

High levels of Mps1 expression are correlated both with the high degrees of aneuploidy and in the aggressiveness a variety of tumor types. Many highly aneuploid tumor cell lines exhibit Mps1-addiction – a dependence on elevated Mps1 expression for their survival ^24–29^. Similarly, when near-diploid HCT116 cells are converted to near-tetraploid cells they too exhibit Mps1 addiction ^30^. It remains unclear, mechanistically, why mammalian cells with abnormally high chromosome numbers have elevated dependence on Mps1. One hypothesis is that having extra-chromosomes places a burden on the chromosome segregation machinery including the demand for Mps1. Previously, it has been shown that cells with increased ploidy are defective in their ability to promote bipolar attachment due to an inadequate upscaling of spindle geometry/size ^31^. But Mps1 has not been evaluated in this model.

To determine if and how cells with increased chromosome numbers become dependent on Mps1, we explored Mps1-dependence in yeast cells with a range of ploidies. We found that polyploid yeast cells, like mammalian cells with elevated chromosome numbers, are addicted to high levels of Mps1 expression. In both haploid and tetraploid yeast, initial erroneous microtubule attachments can be corrected, but the efficiency depends on the spindle length, which grows with ploidy. As ploidy and spindle lengths increase the ability to promote the formation of correct attachments before anaphase onset is greatly compromised, especially when Mps1 levels are reduced.

## RESULTS

### Modulating Mps1 activity

To evaluate whether there are differential requirements for Mps1 activity when ploidy varies, we evaluated chromosome behavior in cells with haploid to tetraploid chromosome content and varied amounts of Mps1 activity. In our initial experiments, Mps1 activity was modulated by combining different copy numbers of the wild-type allele (*MPS1*) or a hypomorphic allele (*mps1-R170S*) ^32^ with copies of the *MPS1* deletion allele (*mps1Δ*). The *mps1-R170S* mutation causes an amino acid change in a domain of the protein defined previously as required for the spindle checkpoint and orienting chromosomes on the spindle^32,33^. This domain is located far from the kinase domain ^34^. Instead, amino acid 170 is positioned at the interface of the Mps1 protein and the Ndc80 complex where Mps1 docks on the outer kinetochore^35,36^. Cells bearing the *mps1-R170S* allele are viable and without noticeable defects in spindle body duplication, which is dependent upon the kinase activity of Mps1^32^. However, *mps1-R170S* mutants exhibit defects in the spindle checkpoint and in meiotic chromosome segregation^32^.

### Dependence on Mps1 activity increases with cell ploidy

To determine whether yeast cells of higher ploidy have an elevated dependence on Mps1, as is seen in tumor cells with elevated chromosome numbers, we varied the ratio of *MPS1* copies/genome in diploid and tetraploid cells (Fig. 1 A). Because low Mps1 activity results in genome instability, cells with reduced *MPS1* copy number in this experiment also carried a plasmid with an additional *MPS1* gene (OPL151). The plasmid carries a centromere, an origin of DNA replication and the *URA3* gene, which allows growth in medium lacking uracil. Previous studies have shown that cells with excess *MPS1* trigger the spindle assemble checkpoint resulting in reduced proliferation ^15^. Because of this, in our experiments (below), which rely on a plasmid shuffle approach, our wildtype control cells carried a centromere plasmid with *URA3* but no *MPS1* gene. Cultures of diploid cells with 0, 1, or 2 chromosomal copies of *MPS1*, and tetraploid cells with 0, 1, 2, 3, or 4 chromosomal copies of *MPS1* were grown selectively in medium without uracil. The cultures were transferred to non-selective medium (YPAD) to allow cells that had lost their *URA3* plasmid to divide and then cells were plated on a medium (5-FOA) that would permit the growth of cells that had lost the plasmid and therefore only had chromosomal copies of *MPS1* (Fig. 1 A). Cells in each culture were subjected to a series of 10-fold dilutions then spotted on media to monitor cell viability. For both diploid and tetraploid cells, complete loss of *MPS1* is lethal (Fig. 1A, 0 copies). The growth of diploid cells with either one or two copies of *MPS1* was indistinguishable. In contrast, for tetraploid cells, viability drops with copy number, most clearly dropping more than tenfold when *MPS1* copy number is reduced from two to one per tetraploid cell.

**Figure 1.**
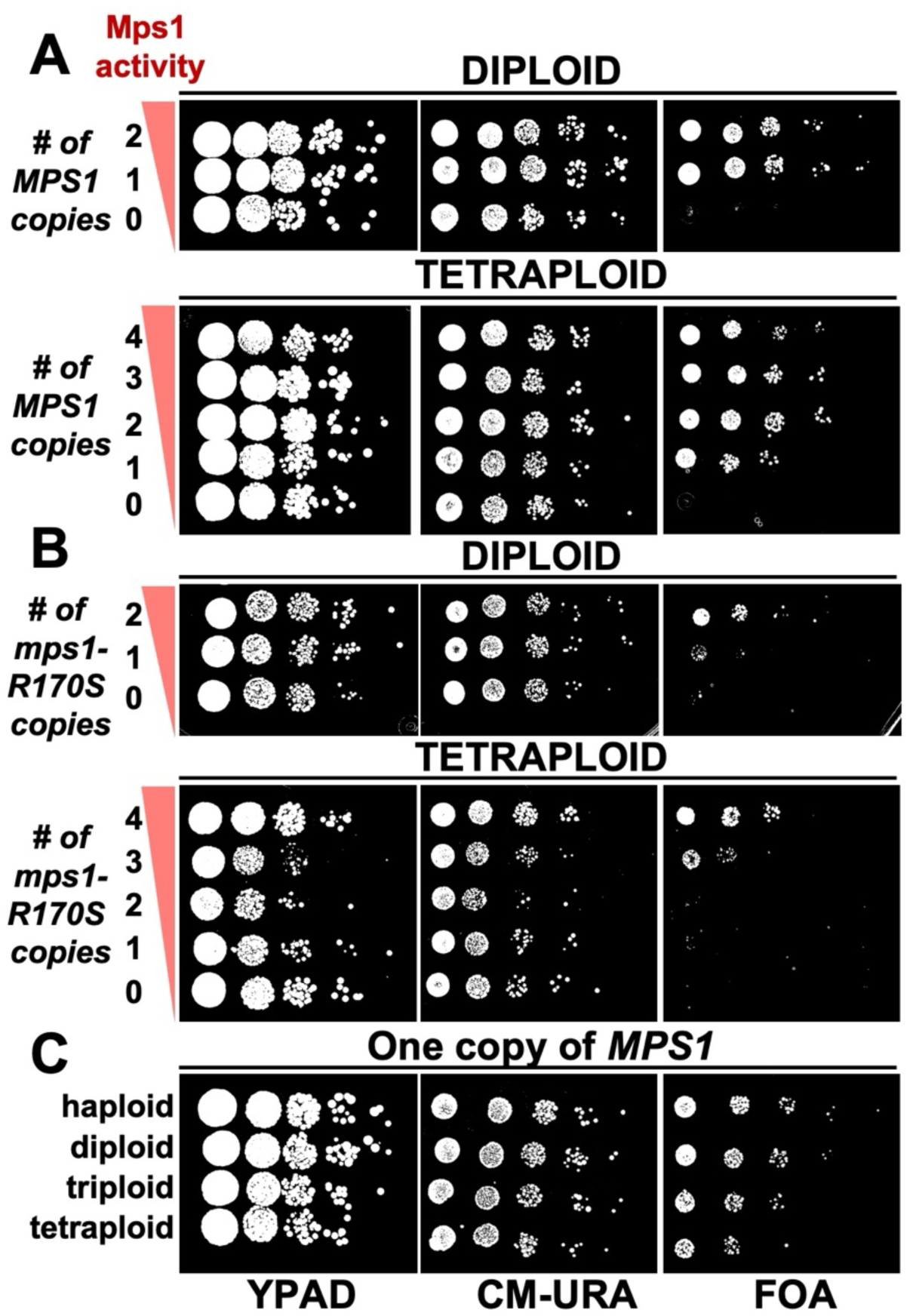
The *mps1* mutants display a ploidy-specific sensitivity. (A-C) Ten-fold serial-dilution growth assays. All strains were propagated carrying a centromere plasmid with the *URA3* selectable marker (either pRS416 with no *MPS1* gene or OPL571 with a wildtype *MPS1* gene). Cells with mutated chromosomal copies of *MPS1* carried OPL571 to support normal cell growth until the time of the experiment. Cells with only wildtype chromosomal *MPS1* genes carried pRS416. Cells were propagated overnight in medium lacking uracil, transferred to rich medium for about two cell generations during which time cells that lose the plasmid can continue to divide, then subjected to ten-fold series dilutions and later spotted on the indicated plates. The FOA plates select for growth of cells that have lost their *URA3* plasmid (and therefore reveals the growth behavior when only the chromosomal set of *MPS1* alleles are available. (A) The growth of diploid cells and tetraploid cells with decreasing numbers of chromosomal *MPS1* copies. (B) The growth of diploid cells and tetraploid cells with decreasing numbers of chromosomal *mps1-R170S* copies. (C) The growth of yeast of different ploidies, each with one chromosomal copy of *MPS1*.

The relationship of ploidy and the need for Mps1 activity is clearer in cells with the hypomorphic *mps1-R170S* allele (Fig. 1 B). Diploid cells show greatly reduced viability with a half dose of *mps1-R170S* per genome (one copy of *mps1-R170S*) while tetraploid cells are completely inviable with a half dose of *mps1-R170S* (two copies). Thus, yeast cells, like mammalian cells, exhibit a ploidy-dependent sensitivity to the levels of Mps1 activity.

This elevated dependence on Mps1 activity as a function of increasing chromosomal content was further examined by comparing the growth of haploid, diploid, triploid, and tetraploid strains each carrying a single copy of *MPS1* (Fig. 1 C). In cells carrying a single copy of *MPS1*, viability drops five-to-tenfold with each increase in ploidy.

### Reducing Mps1 activity in tetraploid cells results in cell death and metaphase delays

Mps1 is essential for duplicating SPBs, the spindle checkpoint, and promoting the proper attachments of chromosomes to the spindle^10^. To determine which among these functions are especially critical for the proliferation polyploid cells, we monitored cellular events using live cell imaging in tetraploid cells expressing varied doses of Mps1 activity. We labeled SPBs with GFP (Spc29-GFP) to facilitate the monitoring of SPB duplication and the timing of metaphase duration (the time from SPB separation to form the metaphase spindle until the rapid extension of spindle length that signals anaphase entry). Cells were placed in microfluidic chambers and imaged for eight hours. To allow propagation of the cells prior to initiating the experiment and allow rapid manipulation of Mps1 activity in the microfluidics chamber, some strains (see below) carried copies of the *MPS1-AID\** allele^37,38^. In these cells Mps1-AID* protein could be quickly eliminated by addition of auxin to the microfluidic chamber prior to initiating the imaging protocol. We initially confirmed that tetraploid cells expressing *MPS1-AID\** (all four copies) were non-viable in the presence of auxin, and that Mps1-AID* is rapidly degraded upon auxin addition (Supl. Fig. 1).

We first tested whether there is a failure of proliferation when the levels of Mps1 are reduced in tetraploid cells. Haploid cells (one copy of *MPS1*), tetraploid cells (four copies of *MPS1*) or tetraploid cells with one functional copy of *MPS1* were monitored for their viability following cell division (Fig. 2 A and Supl. Fig. 2). Haploid cells with one copy of *MPS1* and tetraploid cells with four copies showed low levels of cell death following cell division (4.9% vs. 7.0%). In contrast tetraploid cells with one copy of *MPS1* (MPS1-1X) exhibited nearly 40% death of daughter cells after cell division (Fig. 2 A). This cell death phenotype is not due to reductions in spindle checkpoint activity since it is not seen in tetraploid cells with four copies of *MPS1* but deleted for all four copies of the *MAD2* gene (Fig. 2 A). Similarly, it is not due to failures in SPB duplication since the tetraploid cells with one copy of *MPS1* duplicate their SPBs in nearly every cell division (Supl. Fig. 2).

**Fig 2.**
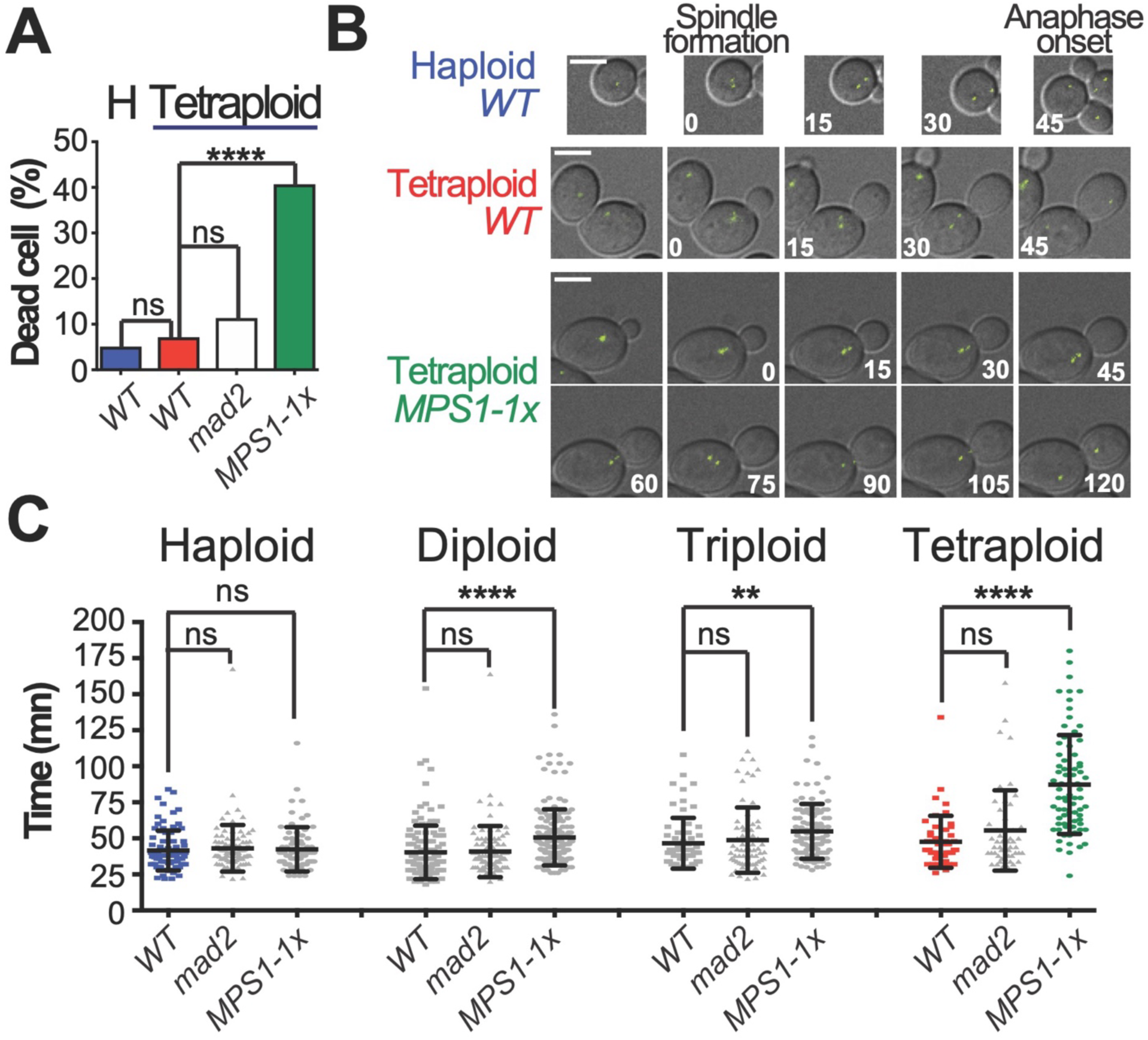
Anaphase onset is delayed in cells with fewer than one *MPS1* gene per genome. (A-C) All cells (haploid, diploid, triploid and tetraploid) are carried a single copy of Spc29-GFP. The cells were grown in defined media and observed by time-lapse imaging at 120-150 second intervals for 4-8 hours. The *MPS1-1x* cells expressed *P_CUP1-_AFB2*, a F-protein necessary for the degradation in yeast cells of the AID* tagged protein. For the *MPS1-1x* genotype, a single *MPS1* allele remains in all ploidies. For the diploid *(MPS1/mps1-AID*)*, triploid *(MPS1/mps1-AID*/mps1Δ)* and tetraploid *(MPS1/mps1-AID*/mps1Δ/mps1Δ),* auxin and copper were added to the media to remove the AID* tagged version of Mps1. (A) The proportion of dead cells was determined during the 1^st^ two cell cycles following each cell division (see Supl. Fig. 2 for details). **** = p < 0.0001, ns=non-significant (Fischer’s exact test). (B) Representative pictures from wild-type cells (haploid and tetraploid) and *MPS1-1x* tetraploid cells are shown. Bars 5μm. Indicated time is in minutes. (C) Time from SPB separation to anaphase onset was measured for each cell of the indicated genotype and ploidy. Strains designated *WT* had one copy of *MPS1* per genome (haploid, *MPS1*; diploid *MPS1/MPS1*; etc.). Cells designated *mad2* were deleted for all copies of the gene. Strains designated *MPS1-1x* carried a single wild-type copy of *MPS1* with the remaining copies of the gene either a deletion or the *MPS1-AID\** version (to facilitate propagation) and had the following genotypes: haploid (*MPS1*), diploid (*MPS1/mps1-aid\**), triploid (*MPS1/mps1-aid*/mps1Δ*), tetraploid (*MPS1/mps1-aid*/mps1Δ/mps1Δ*). Data are data from at least three independent experiments. Each dot represents one data point collected from time-lapse microscopy. ** = p < 0.01, **** = p < 0.0001, ns=non-significant (two tailed unpaired t test). Colored bars and dots in panels A - C are the same strains. Error bars represent one standard deviation.

### Does reducing *MPS1* dosage trigger metaphase delays?

The finding that tetraploid cells with low levels of Mps1 are proficient in SPB duplication and experience cell death that is not likely due to deficiencies in the spindle checkpoint suggest that the vulnerability of the tetraploid cells to *MPS1* dosage is due to other roles of Mps1. Prior studies have shown that Bub1 and Sgo1 are among a group of proteins that are essential for tetraploid cells but not cells of lower ploidy^31^. Bub1 is a component of the spindle checkpoint but has spindle checkpoint-independent roles in chromosome bi-orientation^39,40^. Similarly, Sgo1 protects centromeric cohesion but also has roles that impact bi-orientation^40–42^. It is these bi-orientation functions of Bub1 and Sgo1 that appear to be critical for the survival of tetraploid cells ^42^. Mps1 mediates localization of Bub1 and Sgo1 to the kinetochores and has been implicated in being critical to tetraploid cell survival by impacting chromosome biorientation^17,42,43^. To address this possibility, we tested whether polyploid cells with reduced Mps1 activity experience metaphase delays that might be consistent with a bi-orientation defect. We monitored metaphase duration in haploid, diploid, triploid, and tetraploid cells with either a full complement of *MPS1* genes or a single functional copy per cell (fig. 2 B). Cells were propagated in microfluidic chambers and the time from spindle formation until anaphase onset was measured (Fig. 2 B). In wild-type cells, there is a small increase in metaphase duration with increasing ploidy (Fig 2 C; haploid vs triploid, 41.5 min vs. 46.6 min p=0.049; haploid vs tetraploid 41.5 min vs 47.6 min, p=0.029; unpaired t tests). This increase is consistent with the hypothesis that polyploid cells are confronted with bi-orienting larger numbers of chromosomes and this takes more time. This increase in metaphase duration might be expected to be due to spindle checkpoint-mediated delays as ploidy increases. However, deleting *MAD2* did not shorten metaphase duration for the otherwise wildtype strains, regardless of ploidy (Fig. 2 C). This suggests either that the longer metaphases in polyploid cells are not due to checkpoint delays, or alternatively, *MAD2* has other functions that when removed, slow down metaphase.

In contrast, reducing Mps1 levels significantly increased the duration of metaphase in a ploidy dependent fashion (Fig. 2 C). In diploid cells, removing half of the Mps1 led to a ten-minute increase in metaphase duration (*WT* vs *MPS1-1x*, 50.7 min vs. 40.3 min). We observed a similar increase following the removal of two *MPS1* copies in the triploid cells (*WT* vs *MPS1-1x*, 46.6 min vs 54.9 min). Finally, removing three of the four *MPS1* copies in tetraploid cells doubled the duration of mitosis (*WT* vs *MPS1-1x*, 47.6 min vs 87.3 min). Thus, cells with high chromosome numbers are critically dependent on increased Mps1 dosage for efficient progression through metaphase and this dependence is not mainly due to Mps1’s role in the spindle checkpoint.

### Cells with extended metaphases develop longer spindles

As yeast cells bi-orient their chromosomes in pro-metaphase, the spindle gradually gets longer ^6,44–46^. Despite outward forces that push the spindle poles away from each other, spindle is restrained from extending because the bi-oriented sister chromatids are held together by cohesins. The gradual spindle lengthening in pro-metaphase could be due to “cohesion fatigue” the gradual loss of centromeric sister chromatid cohesion ^47,48^. Previous studies suggested that spindle length is not affected by the changes of ploidy ^31^. But this conclusion was drawn by measuring the length of fixed metaphase spindles. By tracking the distance between the SPBs in individual living cells, we were able to monitor spindle length as a function of time spent time pro-metaphase. Example traces from four representative haploid and tetraploid individual cells (Fig. 3 A) reveal that spindle length gradually increases in pro-metaphase until the dramatic increase in length that marks anaphase entry. An analysis of this type with large sets of haploid and tetraploid cells shows that for both haploids and diploids, spindle length increases over time (Fig 3 B). Because tetraploid cells spend longer in prometaphase than haploid cells (Fig. 2) it may be that ploidy doesn’t directly impact spindle length, but rather indirectly increases spindle length by increasing the duration of pro-metaphase. To explore this, we measured the lengths of nascent spindles (the first imaging frame when we detect two SPB signals) and the final prometaphase spindle (pre-anaphase) in the last imaging frame before anaphase in haploid, tetraploid, and tetraploid cells with only one copy of *MPS1* (Fig. 3 C). The results reveal that when spindles form their lengths are not significantly different across ploidies, as was suggested previously ^31^, and are not impacted by *MPS1* dosage (haploid *WT*: 1.40 μm, tetraploid *WT*: 1.38 μm; tetraploid *MPS1-1x*: 1.40 μm). However, by the time of anaphase onset (pre-anaphase, Fig. 3 D), cells with higher ploidies and reduced *MPS1* dosage have significantly longer spindles. Thus, ploidy, per se, does not impact spindle length, but, since cells with higher ploidy spend extended time in prometaphase, they have significantly longer spindles than cells with lower ploidy when they are completing the bi-orientation process.

**Fig 3.**
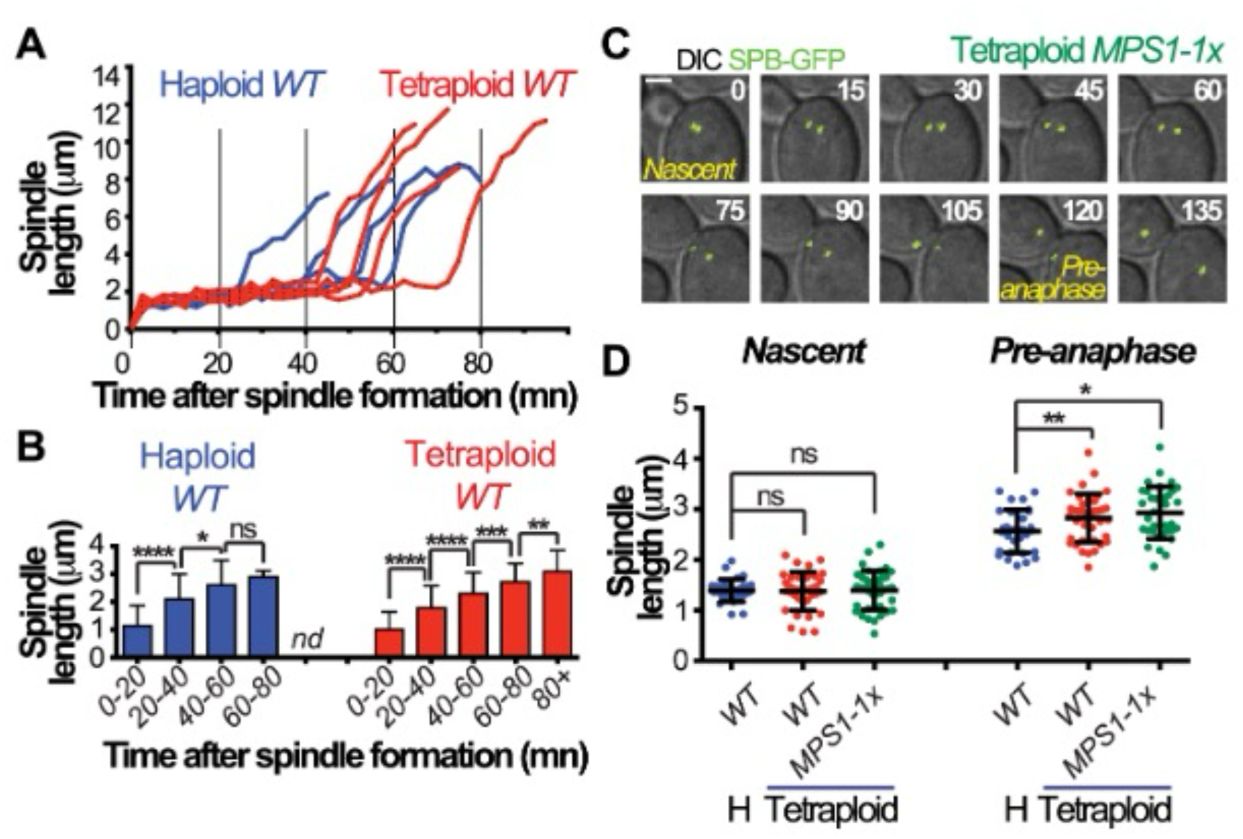
Pre-anaphase spindles get longer when anaphase onset is delay. (A-D) All cells (haploid, diploid, triploid and tetraploid) are carrying a single copy of Spc29-GFP. The cells were grown in SD media and observed by a time-lapse movie at 120-150 second intervals for 4-8 hours. Strain genotypes are as in Fig 2. The spindle length was estimated by measuring in 3D distance between the two SPBs. (A) For each wild-type cell (Haploid and Tetraploid, n≥10), the measurement was done every 150 seconds from prometaphase to the anaphase onset. (B) The graph represents the average of those measurement for the indicated period spent in prometaphase. * = p < 0.05, ** = p < 0.01, *** = p < 0.001, **** = p < 0.0001, ns=non-significant (Unpaired t test). Error bars represent one standard deviation. (C) Representative pictures from the spindle formation (Nascent spindle) to the anaphase onset in a *MPS1-1x* mutant tetraploid cell are shown. Bars 2μm. Indicated time is in minutes. (D) For the indicated genotype, the measurement was either done on nascent spindle or just before the spindle elongate (Pre-anaphase spindle). At least 30 cells were counted per genotype. * = p < 0.05, ** = p < 0.01, ns=non-significant (Unpaired t test). Error bars represent one standard deviation.

### Is bi-orientation efficiency impacted by ploidy or spindle length?

The higher death rate and prolonged metaphases of polyploid cells compared to haploid and diploid cells are consistent with a deficiency in bi-orienting sister chromatids in prometaphase. Two features of polyploid cells that might impact bi-orientation efficiency are their higher chromosome content and longer spindles. Therefore, we tested whether either of these features reduce bi-orientation efficiency. In budding yeast many initial microtubule attachments to sister chromatid pairs are monopolar. These are then corrected to bipolar attachments in prometaphase^6,44,46,49^. During prometaphase the sister chromatid pairs move back and forth across the spindle as they attach and detach from microtubules until bi-orientation is achieved. Previous studies demonstrated that polyploid metaphase cells exhibit a greater proportion of mono-oriented chromosomes than euploid cells^31^. However, since these experiments were done with fixed cells, it was unclear if this phenotype was due to an initial higher frequency of monopolar attachments or to a deficiency in converting monopolar into bipolar attachments. To distinguish between these possibilities, we used live-cell imaging to track chromosome bi-orientation through prometaphase in haploid, diploid, and tetraploid cells. To reveal the movements of chromosomes at higher resolution than in previous experiments, we imaged with faster acquisition rates (5 to 10 second intervals) over the course of five minutes, capturing a window of chromosome behavior in prometaphase. To reduce acquisition times, the SPBs and the centromere of chromosome I (*CEN1*) were both tagged with GFP. We first measured the percentage of cells with that had achieved bipolar attachments of the chromosome I sister chromatids (a centered CEN1-GFP signal) at the beginning of our image acquisition (Fig. 4 A). Cells with short spindles had low frequencies of bipolar attachments and the frequency increased with spindle length. Since our previous experiments showed that spindle length is correlated with the time spent in prometaphase (Fig. 3), we inferred that the cells with very short spindles have just entered prometaphase. The results show that haploid, diploid, and tetraploid cells enter prometaphase with approximately equivalent proportions of monopolar attachments, but they are converted to bipolar attachments more slowly with increasing ploidy (Fig. 4 A). Thus, polyploid cells do not appear to have highly elevated levels of initial monopolar attachments.

**Fig 4.**
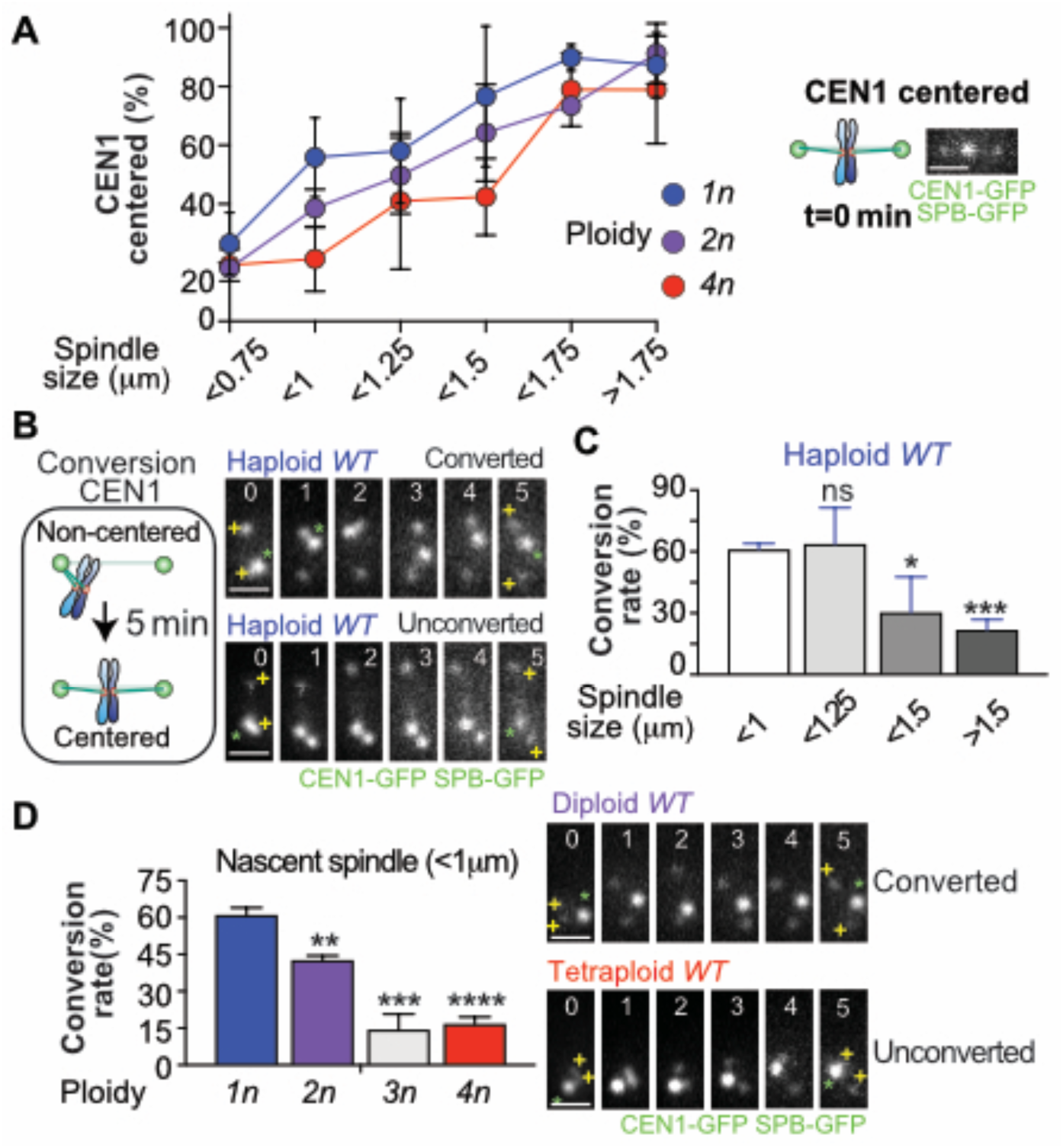
Conversion from monopolar to bipolar attachment is reduced on longer spindles and with higher ploidy. (A-D) All wild-type cells (haploid, diploid and tetraploid) are carrying a single copy of Spc29-GFP and one *CEN1-GFP* tagged chromosome. Cells were observed by time-lapse imaging at 5-10 second intervals for 5 minutes. (A) Cells were placed in categories according to the behavior of *CEN1* at the beginning of the acquisition time (t=0mn). Cells with *CEN1* staying near-by the poles were classified as non-centered (aka syntelic/mono-oriented). Cells with *CEN1* staying in the middle of the spindle were classified as centered (aka amphitelic/bi-oriented) with either un-separated or separated sister chromatid (if their splitting was observed). The proportion of cells in each category, as defined by spindle length, was estimated for the indicated ploidy. At least 137 cells were counted per ploidy. ** = p < 0.01, ns=non-significant (Fischer’s exact t test). Representative pictures of the type of attachment are shown. Bars 1μm. (B) The ability of individual cell to convert monopolar to bipolar attachment in 5 minutes was evaluated. Representative pictures of cells able or unable to convert monopolar attachment are shown. The green asterisks represent CEN1 and the yellow plus signs represent the SPBs. Bars 1μm. Indicated time is in minutes. (C-D) The proportion of cells able to convert in 5mn interval depending on the spindle size in haploid wild-type cells (C) or the indicated ploidy on nascent spindle (D) are shown. At least 20 cells per category of size or ploidy were counted. * = p < 0.05, **** = p < 0.0001, ns=non-significant (Fischer’s exact t test). Representative pictures of cells with nascent spindle able or unable to convert monopolar to bipolar attachment are shown. The green asterisks represent *CEN1* and the yellow plus signs represent the SPBs. Bars 1μm. Indicated time is in minutes.

To test whether polyploid cells are inefficient at converting monopolar to bipolar attachments, we developed an assay to measure conversion efficiency. We assigned cells into one of three categories based upon the position of CEN1-GFP on the spindle. These included monopolar attachment (sister centromeres together and on one side of the spindle (within 0.5 µm of a SPB), bipolar attachment with un-separated sister centromeres (revealed by a central position of the CEN1-GFP focus on the spindle), or bipolar attachment with separated sister centromeres (two CEN1-GFP foci). We then analyzed the ability of cells to convert monopolar attachments into bipolar attachments in a five-minute imaging window (Fig. 4 B). Conversion frequency was then plotted as a function of spindle length. Among haploid cells, those with short spindles were the most efficient while those with longer spindles exhibited reduced rates of monopolar-to-bipolar conversion (Fig 4 C). This result is consistent with a reduced ability of cells to bi-orient chromosomes as the spindle becomes longer. We then compared monopolar-to-bipolar conversion efficiencies as a function of ploidy in cells with similarly-sized short spindles (Fig. 4 D). We found that increasing the ploidy dramatically reduces the efficiency monopolar-to-bipolar conversion (Fig. 4 D). This very low conversion rate in tetraploid cells with short spindles is not reduced further when the spindle length increases (unlike haploid cells) (Supl. Fig. 4).

### MPS1 dosage in tetraploid is critical for monopolar-to-bipolar conversion

The elevated cell death and prolonged metaphases seen in tetraploid cells with reduced *MPS1* dosage (Fig. 2) is consistent with the model that high Mps1 activity is critical for efficient chromosome bi-orientation in polyploid cells. We tested this using the same monopolar-to-bipolar conversion assays described above (Fig. 4). We reduced Mps1 activity by either expressing a single functional *MPS1* locus (*MPS1-1x*) or two *mps1-R170S* loci (*mps1-R170S-2x*) in tetraploid cells. Both tetraploid strains with reduced Mps1 activity exhibited an increased frequency of cells with monopolar attachments on the nascent spindles (Fig. 5 A). Further, the level of bipolar attachments in cells with longer spindles never reached the levels seen in WT tetraploid cells (Fig. 5 A). The result is explained by the finding that both strains with reduced Mps1 activity had lower monopolar-to-bipolar conversion efficiencies (Fig. 5 B). To study the impact of this lower rate of conversion on the final attachments of sister chromatids at anaphase, we monitored the configuration of CEN1-GFP on the spindle in the minute before anaphase entry (t=-60 seconds, pre-anaphase spindle, Fig. 5 C). In haploid cells, just before anaphase (t=-60 seconds) the sister centromeres are usually stretched apart (>90% of cells) on the spindle (Fig. 5 D). In WT tetraploid cells there is a significant reduction in cells with this configuration – instead, the CEN1-GFP was often centered on the spindle but less stretched apart (Fig. 5 D). In contrast, *MPS1-1x* tetraploid cells exhibited a high proportion of cells with non-centered attachments at this stage (Fig. 5 D). Most of these non-centered attachments were probably bipolar, as only one in fifteen cells of the cells scored in this category (Fig. 5 D, *MPS1-1x*, non-centered) non-disjoined when anaphase ensued. Thus, polyploidy itself reduces monopolar-to-bipolar conversion frequency, and this is exacerbated if Mps1 activity is not increased commensurate with the increases in chromosome number.

**Figure 5.**
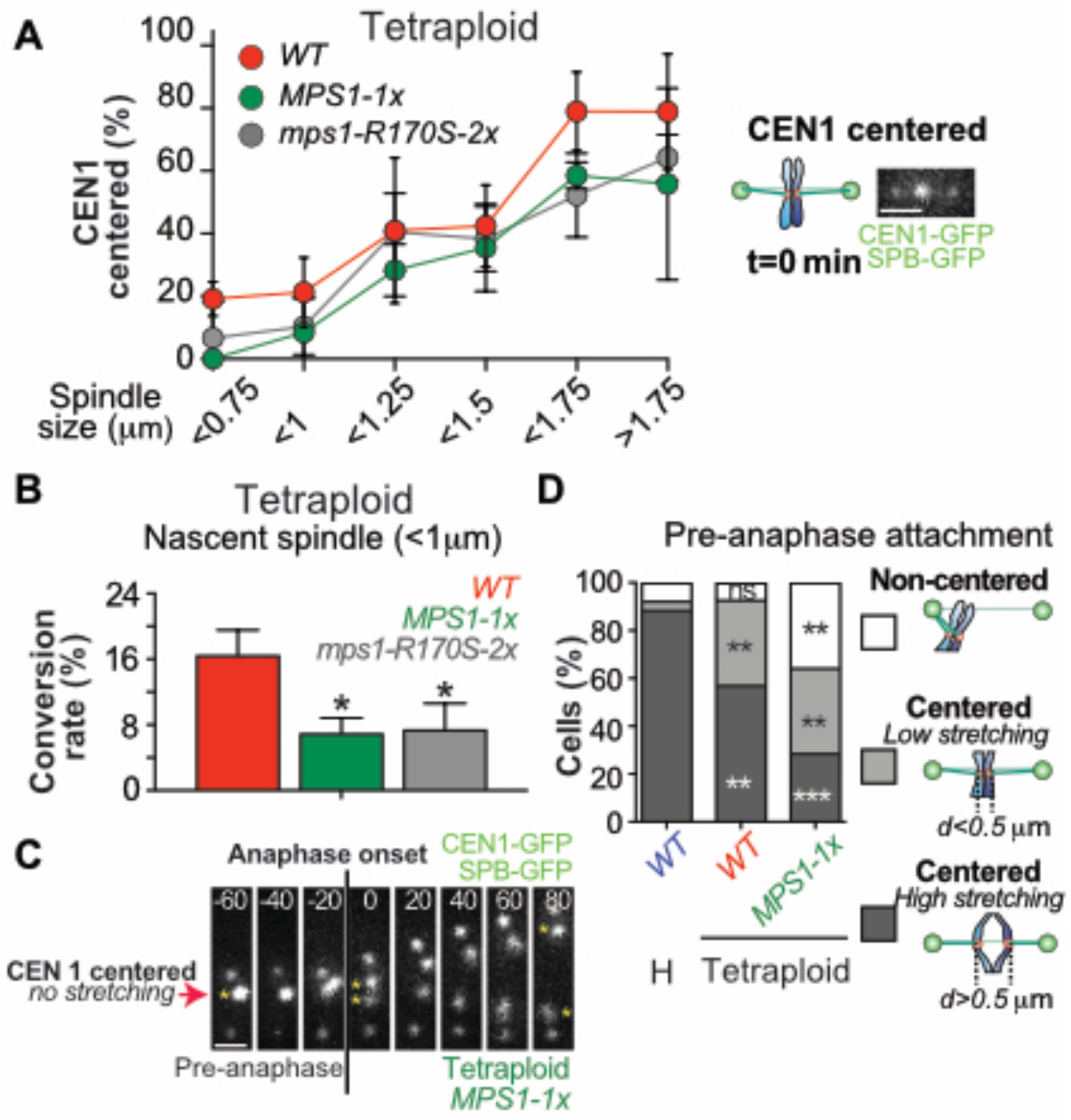
The formation of bipolar attachment is delayed in *mps1* mutants. (A-D) All wild-type and *MPS1-1x* mutant cells (haploid and tetraploid) are carrying a single copy of Spc29-GFP and one *CEN1-GFP* tagged chromosome. Cells were observed by time-lapse imaging at 5-10 second intervals for 5 minutes. (A-B) The assays are the same as described in Fig 4. (A) The proportion of each category, based on the indicated spindle length, is shown for the specified genotype of tetraploid cells. At least 170 cells were counted per genotype. * = p < 0.05, ** = p < 0.01, ns=non-significant (Fischer’s exact t test). (B) The conversion rate of tetraploid cells of the indicated ploidy on nascent spindle (<1μm) was estimated. At least 39 cells per genotype were counted. (C-D) The type of attachment in haploid or tetraploid cells of the indicated genotype was evaluated 1 minute before the anaphase onset. Cells were placed in categories according to *CEN1* behavior at this time (t=-60s). Cells with *CEN1* staying near-by the poles were classified as monopolar attachment. Cells with *CEN1* staying in the middle of the spindle were classified as bipolar attachment with either low stretching (if the distance between the two sister chromatids was lower than 0.5μm) or high stretching (if the distance between the two sister chromatids was higher than 0.5μm). (C) Representative pictures of *MPS1-1x* mutant tetraploid cells displaying a bipolar attachment with low stretching one minute before the anaphase onset are shown. Indicated time is in seconds. Bars 1μm. (D) The proportion of each category depending on the indicated ploidy and genotype is shown. At least 14 cells were counted per genotype and ploidy. ** = p < 0.01, *** = p < 0.001, ns=non-significant (Fischer’s exact test).

### Mps1 acts through Bub1 to promote bi-orientation in tetraploid cells

Previous studies have shown that tetraploid cells are sensitive to reductions in Bub1 activity^42^ and studies in haploid and diploid yeasts have shown that one role of Mps1 is to phosphorylate the kinetochore component Spc105 to allow docking of Bub1/Bub3 on the kinetochore^17,43^. Therefore, we tested whether the sensitivity of tetraploid cells to reductions in Mps1 activity are due to reductions in the contributions of Bub1 to bi-orientation. First, in diploid cells we confirmed that reducing Mps1 activity resulted in a failure in Bub1-GFP recruitment to metaphase kinetochores (Spc24-mCherry) (Fig. 6 A) as described previously^17,43^. Even in diploids expressing a single copy of *mps1-R170S*, there was a significant drop in detectable localization of Bub1 to metaphase kinetochores, commensurate with decreased localization of Mps1 to kinetochores when the R170S region of Mps1 is mutated ^35,50^. If reductions in Bub1 loading at kinetochores is the major reason that tetraploid cells are vulnerable to low Mps1 activity, then mutations that eliminate Bub1 loading at kinetochores should be more detrimental to the proliferation of tetraploids than haploids. To test this, we used viability/growth assays to compare haploid and tetraploid proliferation when Bub1 activity at kinetochore is abolished (Fig. 6 C). In haploid cells, eliminating Bub1 (*BUB1-AID\** plus auxin) has a slight impact on growth (The lethality of the *MPS1-AID\** on auxin containing medium auxin shows that the system is functional). Similarly, haploid cells expressing only a version of Spc105 that cannot be phosphorylated by Mps1 (Spc105-6A) show only partially reduced growth. In contrast, in tetraploid cells, eliminating Bub1 or expressing only the *spc105-6A* allele profoundly diminishes cell viability. If the critical function of *MPS1* in tetraploid cells is converting monopolar-to-bipolar spindle attachments thru Bub1, then the conversion efficiency should be reduced in tetraploid cells when Bub1 activity is reduced. Like tetraploid cells with low dosages of Mps1, cells with no Bub1 show low levels of bipolar attachments of CEN1-GFP upon entry into prometaphase and never reach WT levels of bipolar attached *CEN1* (Fig. 6 D). Indeed, on new spindles, tetraploid cells with depleted Bub1 levels exhibit a significant reduction in the rate at which monopolar attachments are corrected (Fig. 6 E).

**Figure 6.**
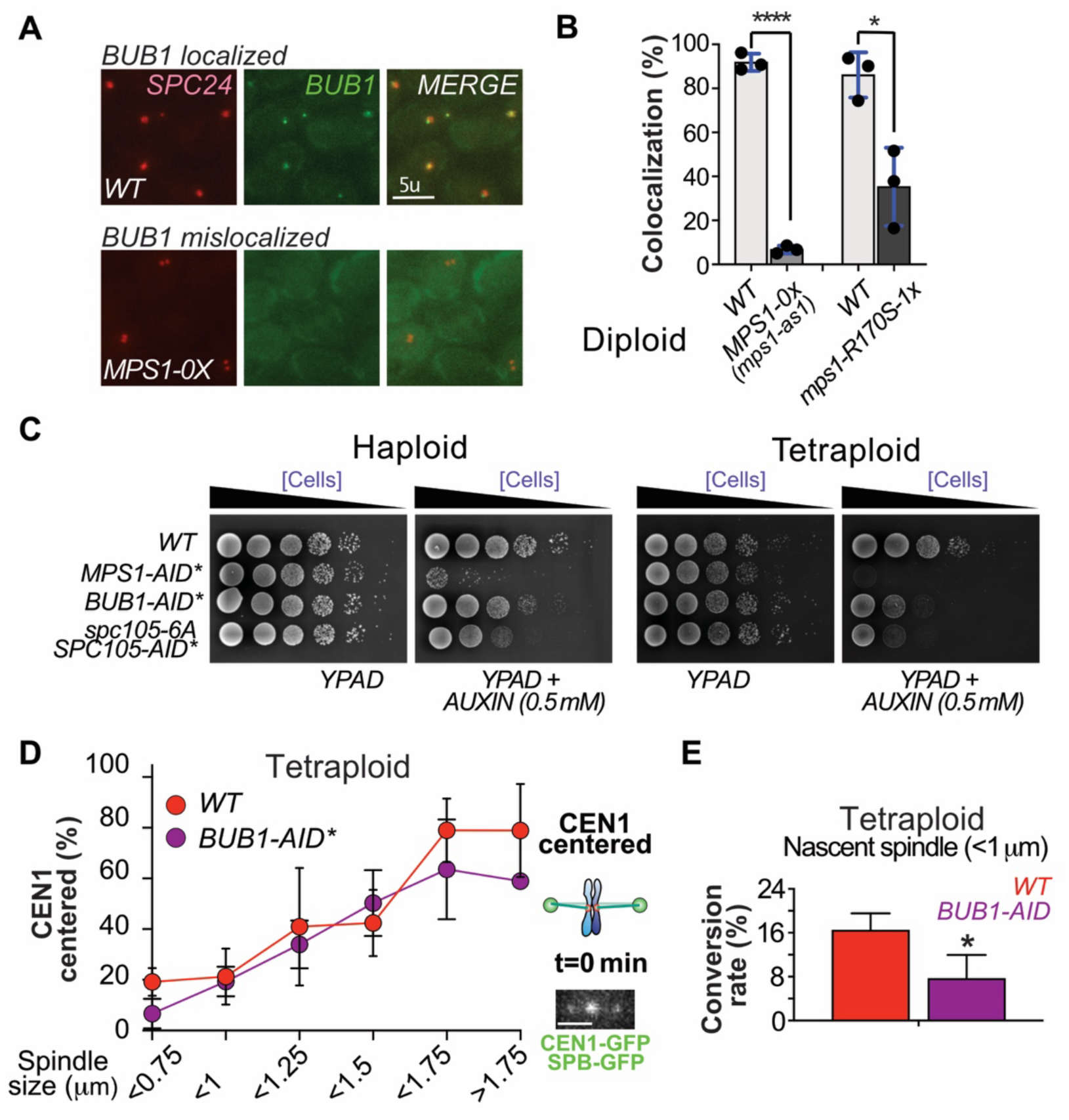
Mps1 acts through Bub1 to promote biorientation in tetraploid cells. (A-B) Diploid cells expressing Bub1-GFP and Spc24-mCherry (to mark kinetochores) were arrested in metaphase (CDC20-AID* plus auxin). MPS1-0x diploids expressed two copies of *mps1-as1*. 1-NM-PP1 was added to the medium to inactive Mps1-as1. *msp1-R170S* diploids expressed one copy of *mps1-R170S* and one copy of *mps1-as1* whose product was inactivated by 1-NM-PP1. The percent of Spc24-mCherry foci with an overlapping Bub1-GFP focus were scored for at least 43 cells in three independent replicates. (C) Haploid cells expressed: one copy of *MPS1*, *MPS1-AID\**, *BUB1-AID\**, or one copy of *spc105-6A* plus one copy of *SPC105-AID\**. Tetraploid cells expressed four copies of *MPS1*, four copies of *MPS1-AID\**, four copies of *BUB1-AID\** or four copies *spc105-6A* plus four copies of *SPC105-AID\**. Cells were grown on permissive conditions then 10-fold dilutions were spotted onto YPAD plates with or without auxin. (D-E) Tetraploid strains (WT or with four copies of *BUB1-AID\**) were imaged (five minutes) to monitor positions of CEN1-GFP between Spc29-GFP marked spindle poles. (D) The percent cells with centered CEN1-GFP as a function of spindle length are shown (from three independent replicates are shown). (E) The percent of cells with nascent spindles in which CEN1-GFP converted from non-centered to centered are shown (* = p < 0.05, Fischer’s exact test).

## DISCUSSION

Polyploid cells are compromised in their fidelity of chromosome segregation, probably because the chromosome segregation apparatus is not adapted to the elevated numbers of chromosomes that must be distributed. In support of this idea, yeast cells with increased ploidy are particularly sensitive to the inactivation of key actors of the chromosome segregation apparatus ^31,42^. Cells with increased ploidy were shown to display more erroneous (= monopolar) attachments on the metaphase spindle than haploid cells, the basis for this phenotype was unclear.

To address this, we tracked chromosome re-orientation as a function of both ploidy and Mps1 levels using live-cell imaging. The efficiency with which euploid and polyploid cells initially attach chromosomes correctly on the spindle was indistinguishable. For both euploid and polyploid cells most initial attachments of sister chromatids to the spindle are monopolar – probably because the kinetochores of most of the spindles are clustered near one pole when the spindle forms ^6^. The impact of increasing chromosome numbers is that as ploidy increases cells exhibit a diminished ability to convert monopolar attachments to bipolar attachments. Here we identified two factors that impact the efficiency of correcting monopolar attachments. The first is spindle length. In haploid cells monopolar attachments are rapidly converted into bipolar attachments early in prometaphase when cells have short spindles. But as spindle length increases this ability is greatly reduced. The same is true across all ploidies. Notably however, the duration of prometaphase increases with ploidy, and since spindle length gradually but continuously increases during prometaphase, cells with very long spindles are more common among polyploid cells. A second factor that impacts the efficiency with which cells bi-orient their chromosomes is chromosome content. Cells with more chromosomes convert monopolar attachments into bipolar attachments less efficiently than cells with fewer chromosomes, even when they have the same spindle length. It is unclear why this is so. The number of kinetochores and microtubules increase proportionately with ploidy^31^, but it seems likely that some other critical factors do not scale when ploidy increases.

To find why cells are so dependent on Mps1 when the chromosome number increases, we reduced Mps1 levels and studied its impact on chromosome segregation. We identified four striking features in *mps1* mutant cells: 1) The transition from metaphase to anaphase is delayed. 2) The spindle length is consequently extended in late prometaphase because of the delay. 3) The conversion efficiency from monopolar to bipolar attachment is reduced. 4) Cells show an increased sensitivity to reduced Mps1 dosage at higher ploidy. Studies in fission yeast have also identified an impact on re-orienting chromosomes when spindle length is increased ^51^. Here, two processes were proposed to act in parallel to move mono-oriented kinetochores from the pole to the spindle mid-zone where chances for achieving a bioriented attachment is increased. These were termed the direct and indirect routes. In the direct route, microtubules from one pole reach across the spindle to connect to mono-oriented kinetochore and drag them back towards the mid-zone (Fig. 7). In the indirect route, also referred to as chromosome gliding, kinetochores were proposed to slide along microtubules that emanate from the pole where they are clustered, toward the plus ends. Such indirect movements have been reported in mammalian cells and fission yeast ^52,53^. In fission yeast, the spindle checkpoint proteins Bub1, Bub3, and Mad1 have been implicated in promoting these movements^52,54^

**Figure 7.**
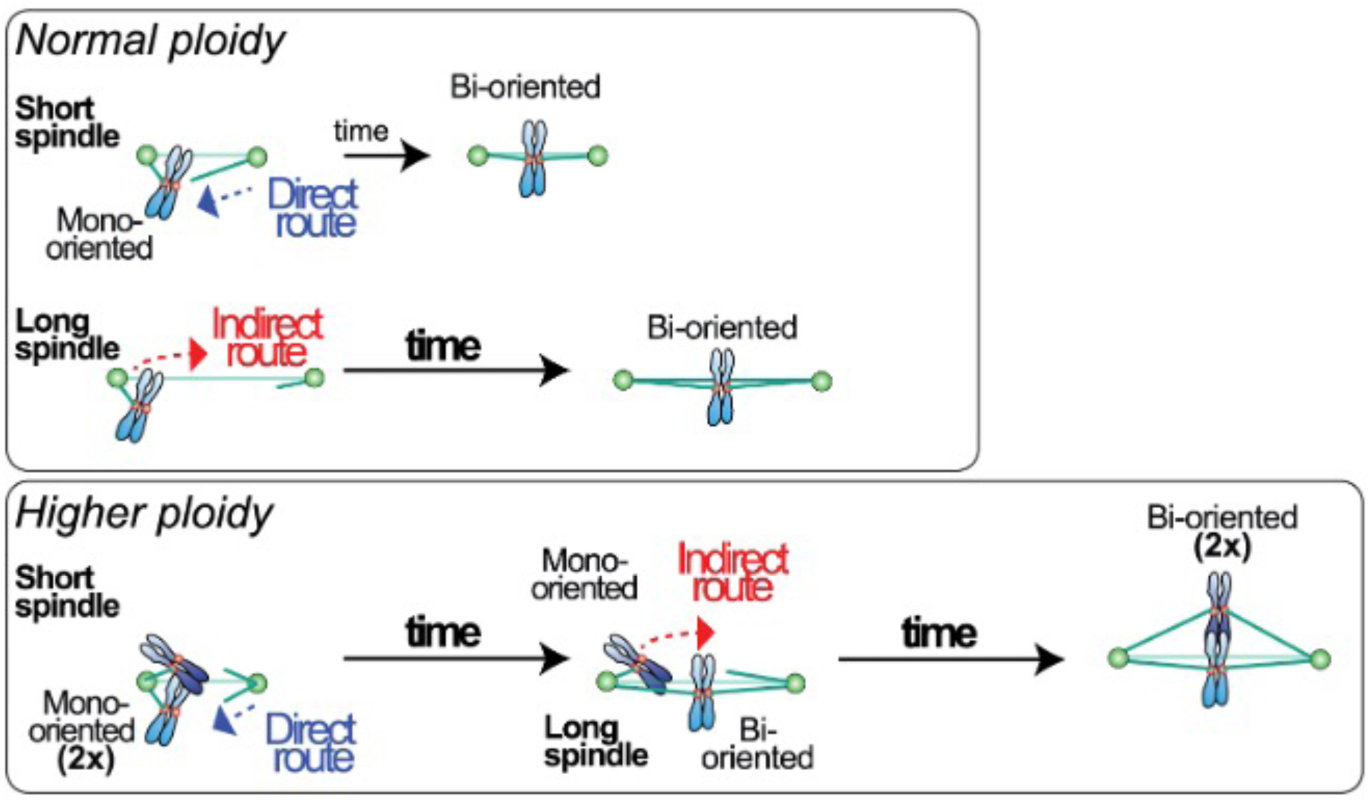
Cells with elevated ploidy may have elevated dependence on alternate pathways for moving chromosomes on the spindle. Cells with normal ploidy can quickly bi-orient their chromosomes. Microtubules from one pole can reach across the spindle to drag chromosomes toward the midzone (direct route). In polyploid cells (2x), the increase in chromosome number slows initial bi-orientation and the spindle elongates in the prolonged prometaphase. On longer spindles the direct route is less efficient and cells depend on an alternate, indirect route, to move chromosomes along the lateral surfaces of microtubules towards the spindle mid-zone

Our data are consistent with the model that Mps1 supports the direct and/or indirect routes of achieving biorientation. Prior work has shown that Mps1 promotes movements of meiotic chromosomes across the spindle that appear to be similar to the direct route – mediated by microtubules that reach from one pole across the spindle to attach to a kinetochore on the other half of the spindle - and moving the chromosome towards the spindle midzone^32,55^ Because the microtubule has to stretch across the spindle, this process would presumably be more efficient in cells with short spindles. However, on longer spindles like those found in polyploid cells with prolonged prometaphases, the direct route may be limited, and chromosomes may rely more heavily on an indirect route. The localization of Bub1 and other spindle checkpoint proteins (Mad1) to kinetochores, a dedicated role of Mps1, has been proposed to be critical for the indirect route^51,52^. Our findings and work by others have shown that polyploid cells are not especially vulnerable to loss of the spindle checkpoint but are sensitive to reductions of Mps1 and Bub1 activity^31,42^. Thus, the vulnerability of polyploid cells to reductions in Mps1 function may be due to an elevated dependence on pathways for chromosome movement that can efficiently help chromosomes to bi-orient on spindles with less-than-ideal geometries. Elucidating the mechanistic details of how Mps1 helps chromosomes become bi-oriented may assist in revealing, in more detail, the addiction of highly aneuploid tumor cells to high levels of Mps1 activity.

## MATERIALS AND METHODS

### Yeast strains and culture conditions

All strains are derivatives of two strains termed X and Y described previously ^56^. We used standard yeast culture methods ^57^. The yeast strains used in this study are lister in Table S1-3. For the long-term time-lapse microscopy, the diploid, triploid and tetraploid *MPS1-1x* mutants, auxin (0.5mM, Sigma Aldrich I5148-10G) and copper (200μM, Sigma Aldrich 451657-10G) were added after cells were loaded into the chamber. For the short-term time-lapse microscopy with tetraploid *MPS1-1x* mutants, auxin (0.5mM, Sigma Aldrich I5148-10G) and copper (200μM) were added 30mn before cells were concentrated on coverslips. For the serial-dilution assay, auxin (0.5mM) was added to YPAD plate.

### Generation of triploid and tetraploid cells

The tetraploid cells were generated by mating diploid strains of opposite mating types (Diploid *Mata*/*Mata* with diploid *Mat*α/*Mat*α). The triploid cells were generated by mating a haploid strain with a diploid strain of the opposite mating type (haploid *Mata* with diploid *Mat*α/*Mat*α or haploid *Mat*α with diploid *Mata*/*Mata*).

The diploid strains *Mata*/*Mata* and *Mat*α/*Mat*α were generated by switching one of the mating types of parent diploid strains *Mata*/*Matα.* The mating-type switch was done via two-step gene replacement strategies. Two different plasmids carrying the *URA3* marker and either the *Matα* allele (pSC9 = OPL233) or the *Mata* allele (pSC11 = OPL234) were used ^58^. The diploid strains *Mata*/*Matα* were originally transformed by inserting the plasmids digested by EcoRI at the mating type locus. After transformation, *URA3+* clones were selected. After growing the clones overnight on non-selective media (YPD), the revertant *ura-* clones were selected by using 5’FOA plate that is toxic for *URA+* cells. Finally, the mating type of diploid *ura-* clones were tested.

### Genetic modifications

Heterozygous *CEN1-GFP* dots: An array of 256 lacO operator sites on plasmid pJN2 was integrated near the *CEN1* locus (coordinates 153583–154854). *lacI-GFP* fusions under the control of *P_CYC1_* was also expressed in this strain to visualize the location of the *lacO* operator.

PCR-based methods were used to create complete deletions of ORFs (*mps1::kanMX, mad2::NAT)* ^59,60^*. mps1-R170S::his5* strains were generated previously ^32^.

### SPC29-GFP

*SPC29* was tagged with GFP at its C-terminal at its original locus using one-step PCR method and the plasmid OPL436 (pFA6a-link-yoEGFP-Kan, http://www.addgene.org/44900).

*mps1-aid\**: *MPS1* gene was fused with an auxin-inducible degron tag at its C-terminal at its original gene locus, using the one-step PCR method, as described previously ^37^. Using two plasmids (pKan–AID*–9myc and pNat–AID*–9myc), we generated two different versions of Mps1 C-terminal tagged: *MPS1-AID*-9myc::NAT* and *MPS1-AID*-9myc::KAN*.

*HphNT1-RS1-CEN3-RS1* and *CEN3-RS2-HIS3MX6: HPH+* and *HIS+* markers were inserted at close proximity to the centromere of chromosome 3 to exclude among the *ura-* revertant the one coming from chromosome loss.

F-Box protein *AFB2*: As an auxin receptor, we used *AFB2*, which promote an enhanced degradation in a yeast-based system compare to classically used *TIR1* ^61^. We placed *AFB2* expression under the control of either the *GPD1* (*P_GPD_-AFB2-LEU2*) or *CUP1* promoter (*P_CUP1_- AFB2-HIS*).

Plasmid shuffle: A plasmid containing a copy of *MPS1-as1* and the *TRP1* yeast marker (OPL77 *=* pRS314-*MPS1-as1,* a gift from Mark Winey) or a plasmid containing MPS1 and the *URA3* gene (OPL571) were added to the cells to keep them alive when having a null allele of *MPS1* or prevent the accumulation of suppressor mutations when having the *mps1-R170S* allele. The shuffling plasmids were counter-selected by placing the cells on media containing the compound 5-Fluoroanthranilic acid (FAA) which selects against Trp^+^ cells ^62^ or 5-Fluoro-ortic acid which selects against Ura^+^ cells^63^.

### Fluorescence microscopy

Long-term time-lapse microscopy. Images were acquired in 2-2.5min intervals for 6–8 h with exposure times of 100-200ms according to fluorescence intensity. Cells were imaged using ONIX Microfluidic Perfusion System from CellAsic with a flow rate of 5 psi (http://www.emdmillipore.com/US/en/life-science-research/cell-culture-systems/cellASIC-live-cell-analysis/microfluidic-plates/). Cells were incubated at 30°C to mid–log phase and loaded into the chamber for haploid, diploid and triploid cells. Tetraploid cells were directly loaded into the chamber. Haploid cells were loaded on Y04C plates whereas diploid, triploid and tetraploid cells were loaded on Y04D or Y04E plates. For haploid, diploid and triploid cells, time-lapse imaging was started 30-60mn after the loading into the chamber and was performed at 30 °C. For tetraploid cells, time-lapse imaging was started 3h30-4h after loading into the chamber and was performed at 30 °C.

For the time-lapse imaging following the behavior of the spindle, a marker for SPBs (*SPC29-GFP)* was used. Images were originally collected with a Nikon Eclipse TE2000-E equipped with the Perfect Focus system, a Roper CoolSNAP HQ2 camera automated stage, an X-cite series 120 illuminator (EXFO) and NIS software. Images were later collected with a Nikon Eclipse Ti2, a Hamamatsu ORCA-Flash4.0 V3 Digital CMOS C13440-20CU camera, a Lumencor SPECTRA X light engine and NIS software.

Time-lapse microscopy was analyzed for cell cycle duration. The formation of the spindle was defined as the time when two separated SPBs can be easily observed. Anaphase onset was defined as the time when a rapid elongation of the spindle was observed. Data were graphed in Prism (GraphPad Software) and the significance was calculated using the Unpaired t test.

High speed time-lapse microscopy. Time-lapse imaging (every 5-10 second for 5-10 minutes) were collected using a Roper CoolSNAP HQ2 camera on a Zeiss Axio Imager 7.1 microscope fitted with a 100×, NA1.4 plan-Apo objective (Carl Zeiss MicroImaging), an X-cite series 120 illuminator (EXFO). Cells were incubated at 30°C to mid–log phase. Dividing cells were concentrated, spread across polyethyleneimine-treated coverslips, then covered with a thin 1% agarose pad to anchor the cells to the coverslip ^64^. The coverslip was then inverted over a silicone rubber gasket attached to a glass slide.

The measurements of the spindle length were all done at the beginning of the movies (t=0 min). For the analysis of type of *CEN1-GFP* attachment on bipolar spindle, the position of the centromeres was estimated at the beginning and at the end (t=5 min) of the movie. The centromeres were defined as monopolar when sister chromatids remain un-separated at close proximity (<0.5 μm) to the SPBs for at least three consecutive frames. Centromeres were considered to be bipolar with un-separated sister chromatids when they remain at a central position on the spindle and the sister chromatids appears as a single dot for at least three consecutive frames. Centromeres were considered to be bipolar separated when the sister chromatids were observed at two separated dots for at least three consecutive frames. Cells were defined as converted when able to switch from a monopolar attachment at t=0 minutes to a bipolar attachment at t=5 minutes. Cells were defined as unconverted when observed as monopolar attachment at t=0 minutes and t=5 minutes. For the analysis of type of *CEN1-GFP* attachment on pre-anaphase spindle, the anaphase onset (t=0 seconds) was initially determined by observing a rapid elongation of the spindle. The pre-anaphase attachment was determined by observing the type of attachment one minute before this event (t=-60 seconds). The centromeres were defined as monopolar when the sister chromatids remain un-separated and at close proximity (<0.5 μm) to the SPBs for at least three consecutive frames. Centromeres were considered to be bipolar with low stretching when the distance between the two sister chromatids was under 0.5 μm. Centromeres were considered to be bipolar with high stretching when the distance between the two sister chromatids was over 0.5 μm.

## ACKNOWLEDGEMENTS

We thank current and past laboratory members for reagents, and discussions of our work. AS, DH, MG, and JH contributed to this project as participants in the OMRF Fleming Scholars Program for undergraduate summer research. This project was supported by NCI grant R03-CA212867 to REM, NIH grant R35-GM126980 to GJG, and NSF grant 2029286 and NIH grant R35-GM152165 to DSD.

## Abbreviations

SPB: spindle pole body

