## Supplementary Material for "Yeast with elevated chromosome numbers are addicted to high levels of Mps1"

**SUPPLEMENTAL MATERIAL**

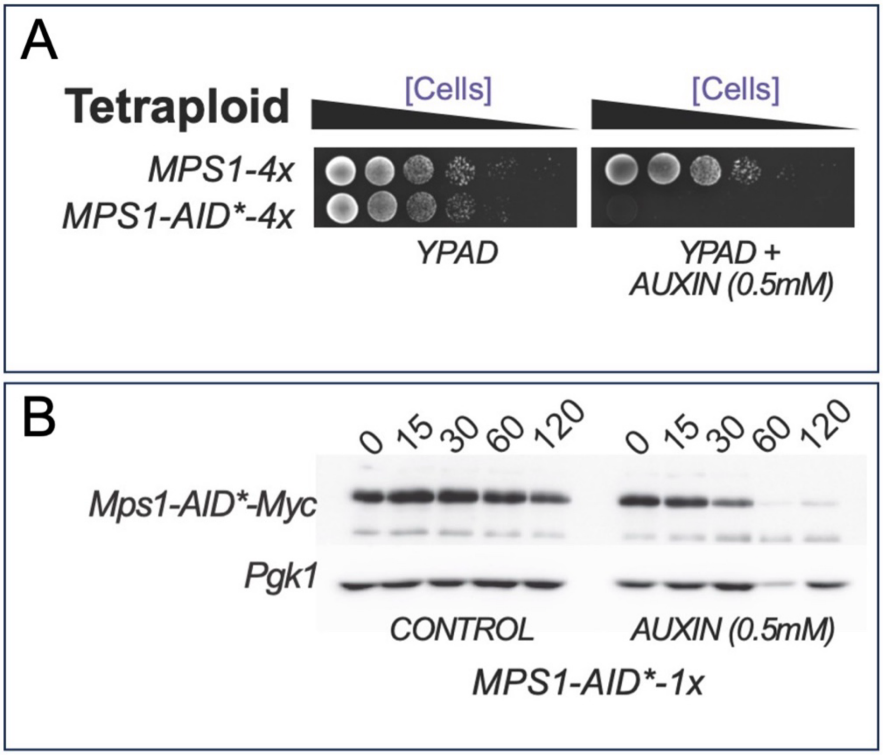

**Supplemental Figure 1. Viability assay on *mps1-AID** tetraploid cells. A.** Ten-fold serial-dilution growth assays for tetraploid cells. Tetraploid cells carrying four *MPS1* loci or four *MPS1-AID** loci and expressing the *AFB2* F-box protein were grown in YPAD medium and then diluted and spotted on YPAD plates with auxin (0.5mM) to induce the degradation of Mps1-AID*. **B.** Western blots showing degradation of Mps1-AID*. Tetraploid yeast expressing a single *MPS1-AID** gene and *AFB2* were grown to log phase in YPAD medium. Auxin (0.5 mM) was added to the culture medium in one flask and cells were harvested at the indicated times. Samples were probed with antibodies versus the MYC epitope and the Pgk1 loading control.

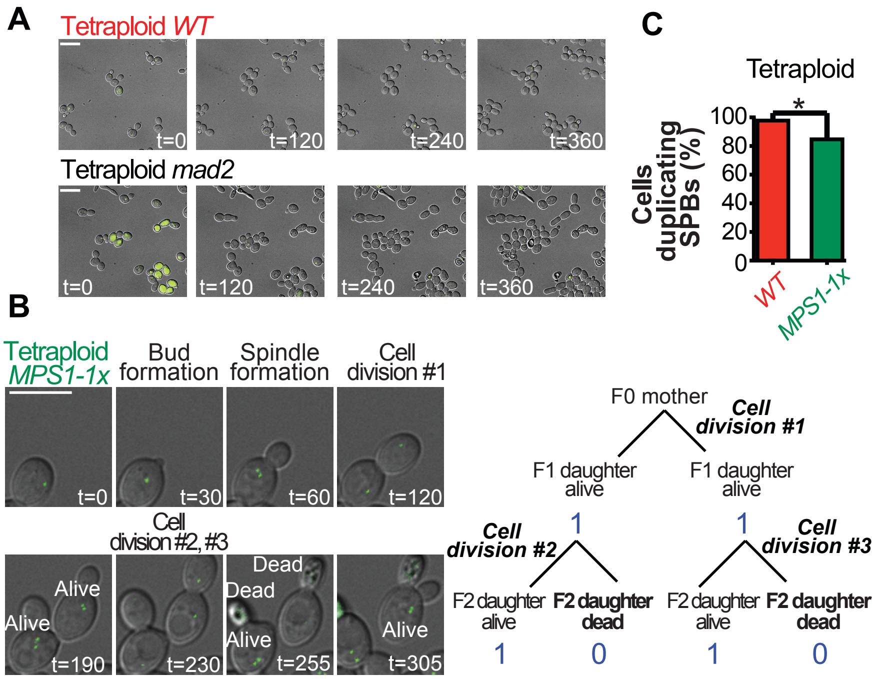

**Supplemental Figure 2. Cell division and SPB duplication assays.** All cells (haploid and tetraploid) are carrying a single copy of Spc29-GFP. (A) Representative pictures of wild-type and *mad2* mutant tetraploid cells growing inside the microfluidic chamber are shown. Indicated time is in minutes. Bars 10µm. (B) The cell viability shown in Fig. 2A was estimated by following the behavior of the progeny during the 1st two cell cycles. Cells scored as "dead" fail to form new buds, and frequently exhibit numerous vacuoles and become auto-fluorescent. We evaluated the proportion of non-growing/dead cells after each cell division. In the example of *MPS1-1x* tetraploid cells, shown, we observed two viable cells after the first cell division (2 of 2). In the second round of cell division (cell division #2 and #3), half of the progeny (2 of 4) were not viable. Indicated times are in minutes. Scale bars =10µm. (C) Are polyploid cells sensitive to Mps1's function in spindle pole body duplication? To identify a potential defect, we evaluated the ability of tetraploid cells to duplicate their SPBs. For this purpose, we used the formation of a bud as a marker the beginning of the cell cycle and tracked whether cells entering the cell cycle were able to display two distinct SPBs. The majority of the *MPS1-1x* tetraploid cells could duplicate their SPBs suggesting this is not the main problem for these cells. However, there was a slight reduction in SPB duplication compared to the *MPS1-4x* tetraploid cells. Our cell death assay (Supl Fig. 2 B) suggests that death usually occurs before entry into a new cell cycle (in G1) but it is possible that it occasionally occurs after cell cycle and before SPB duplication, explaining the slight reduction in SPB duplication in MPS1-1X tetraploid cells (Supl. Fig. 2 C; * = $p < 0.05$, Fischer's exact test).

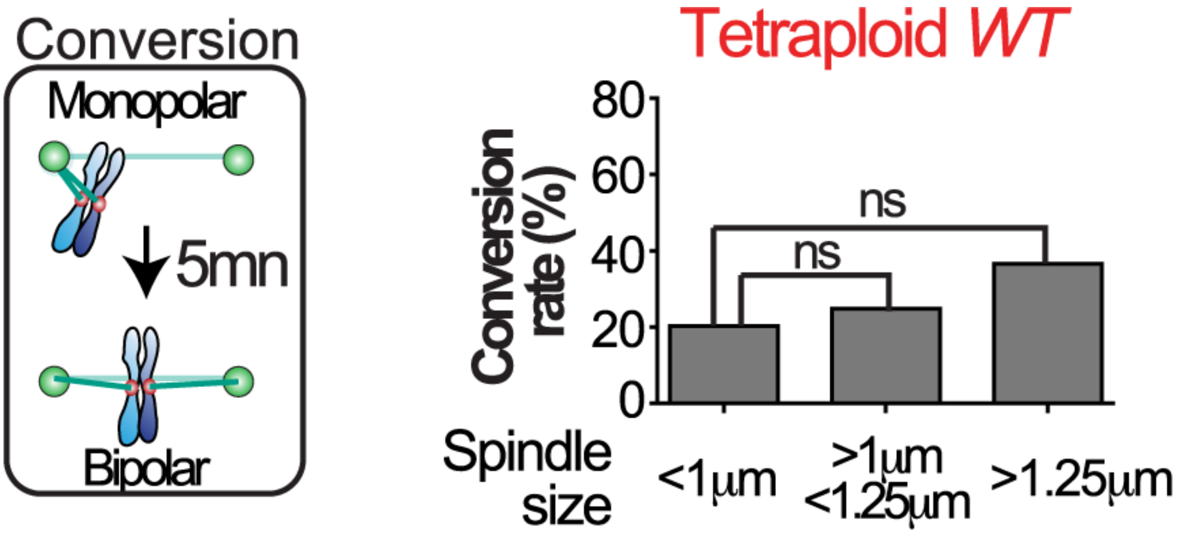

**Supplemental Figure 3. Conversion from monopolar to bipolar attachment in wild-type tetraploid cells.** The ability of individual tetraploid wild-type cells to convert monopolar to bipolar attachment in 5 minutes was evaluated depending of the spindle size. The proportion of cell able to convert in this time interval is shown as a function of spindle length. At least 19 cells were counted in each category. ns=non-significant (Fischer's exact t test).

**Supplemental Table 1. Strains used in each experiment**

| Figure | Ploidy and Genotype | Strains | Parents strain (Haploid or diploid) |
|---|---|---|---|
| **Fig. 1A** | Diploid *WT* | PHC764 | O1195 * O1196 |
| | Diploid *MPS1-1x* | PHC763 | O1195 * O1194 |
| | Diploid *MPS1-0x* | PHC753 | O1193 * O1194 |
| | Tetraploid *WT* | PHC743 | O1191 * O1192 |
| | Tetraploid *MPS1-3x* | PHC744 | O1186 * O1187 |
| | Tetraploid *MPS1-2x* | PHC745 | O1183 * O1188 |
| | Tetraploid *MPS1-1x* | PHC746 | O1183 * O1189 |
| | Tetraploid *MPS1-0x* | PHC747 | O1185 * O1189 |
| **Fig. 1B** | Diploid *mps1-R170S-2x* | PHC517 | O1178 * O1176 |
| | Diploid *mps1-R170S-1x* | PHC518 | O1178 * O1177 |
| | Diploid *mps1-R170S-0x* | PHC511 | O1179 * O1177 |
| | Tetraploid *mps1-R170S-4x* | PHC519 | O1172 * O1173 |
| | Tetraploid *mps1-R170S-3x* | PHC520 | O1172 * O1175 |
| | Tetraploid *mps1-R170S-2x* | PHC521 | O1172 * O1181 |
| | Tetraploid *mps1-R170S-1x* | PHC522 | O1180 * O1174 |
| | Tetraploid *mps1-R170S-0x* | PHC516 | O1180 * O1182 |
| **Fig. 1C** | Haploid *WT* | O1195 | |
| | Diploid *MPS1-1x* | PHC763 | O1195 * O1194 |
| | Triploid *MPS1-1x* | PHC749 | O1184 * O1194 |
| | Tetraploid *MPS1-1x* | PHC748 | O1184 * O1190 |
| **Fig. 2A** | Haploid *WT* | X3177 | |
| **Fig. 2A, 3A-B, S2** | Tetraploid *WT* | 4nDH111<br>4nRM124 | O753 * O630,<br>O750 * O627 |
| | Tetraploid *mad2* | 4nDH113<br>4nRM126 | O849 * O576,<br>O852 * O573 |
| | Tetraploid *MPS1-1x* | 4nDH115<br>4nDH116 | O853 * O860,<br>O856 * O857 |
| **Fig. 2B-C, 3** | Haploid *WT* | X3177, X3178 | |
| **Fig. 2C** | Haploid *mad2* | X3469, X3470 | |
| | Haploid *MPS1-1x* | X3473, X3474 | |
| | Diploid *WT* | O753, O750, O749, O754 | |
| | Diploid *mad2* | O849, O852, O850 | |
| | Diploid *MPS1-1x* | DDH3713, DRM4019<br>DAS4047, DRM4020,<br>DAS4047 | X3473 * X2834<br>X3474 * X2833 |

**Supplemental Table 1. Strains used in each experiment (continued)**

| | | | |
|---|---|---|---|
| **Fig. 2B-C** | Triploid *WT* | 3nDH21<br>3nRM33<br>3nRM34 | O753 * X206<br>O754 * X206<br>O749 * X55 |
| **Fig. 2C** | Triploid *mad2* | 3nDH23, 3nRM35<br>3nDH24 | O849 * X259<br>O852 * X258 |
| **Fig. 2C** | Triploid *MPS1-1x* | 3nDH25, 3nRM37<br>3nRM38 | O853 * X2834<br>O856 * X2833 |
| **Fig. 2B-C, 3D** | Tetraploid *WT* | 4nRM123<br>4nAS135<br>4nAS136 | O753 * O630<br>O754 * O629<br>O750 * O627 |
| **Fig. 2C** | Tetraploid *mad2* | 4nRM125<br>4nAS137<br>4nAS138 | O849 * O576<br>O850 * O575<br>O852 * O573 |
| **Fig. 2B-C** | Tetraploid *MPS1-1x* | 4nRM127<br>4nRM128<br>4nAS140 | O853 * O860<br>O856 * O857<br>O855 * O858 |
| **Fig. 4-5, S3** | Haploid *WT* | X3493, X3494 | |
| | Diploid *WT* | DAS3717, DAS4707 | X3177 * X754 |
| | | DAS3718 | X3178 * X753 |
| | Tetraploid *WT* | 4nAS117 | O753 * O904 |
| | | 4nAS118, 4nAS142 | O750 * O903 |
| | | 4nAS131 | O754 * O904 |
| | | 4nAS132 | O749 * O903 |
| | Tetraploid *MPS1-1x* | 4nAS119, 4nAS143 | O853 * O907 |
| | | 4nAS120 | O856 * O905 |
| | | 4nAS133 | O854 * O908 |
| | | 4nAS134 | O855 * O906 |
| **Fig. 6 A** | WT | DRM4237 | X3799 * X3800 |
| | MPS1-0x | DRM4238 | X3805 * X3806 |
| **Fig. 6 B** | WT | DRM4237 | X3799 * X3800 |
| | MPS1-0x | DRM4238 | X3805 * X3806 |
| | WT | DRM4239 | X3805 * X3804 |
| | mps1-R170S | DRM4250 | X3790 * X3805 |

**Supplemental Table 1. Strains used in each experiment (continued)**

| **Figure** | **Ploidy and Genotype** | **Strains** | **Parents strain** (Haploid or diploid) |
|---|---|---|---|
| **Fig. 6 C** | Haploid WT | X2608 | |
| | Haploid MPS1-AID* | X2837 | |
| | Haploid BUB1-AID* | X3727 | |
| | Haploid spc105-6A/SPC105-AID | X3737 | |
| | Tetraploid WT | O655 | O631 * O633 |
| | Tetraploid MPS1-AID* | O791 | O635 * O639 |
| | Tetraploid BUB1-AID* | 4nAS147 | O924 * O926 |
| | Tetraploid spc105-6A/SPC105-AID | 4nAS145 | O927 * O931 |
| **Fig. 6 D** | Tetraploid WT | 4nAS117 | O753 * O904 |
| | | 4nAS118, 4nAS142 | O750 * O903 |
| | | 4nAS131 | O754 * O904 |
| | | 4nAS132 | O749 * O903 |
| | Tetraploid BUB1-AID* | 4nAS153 | O984 * O926 |
| | | 4nAS154 | O985 * O926 |
| **Fig. 6 E** | Tetraploid WT | 4nAS117 | O753 * O904 |
| | | 4nAS118, 4nAS142 | O750 * O903 |
| | | 4nAS131 | O754 * O904 |
| | | 4nAS132 | O749 * O903 |
| | Tetraploid BUB1-AID* | 4nAS153 | O984 * O926 |
| | | 4nAS154 | O985 * O926 |
| **Fig. S1** | Tetraploid *MPS1-4X* | O655 | O631 * O633 |
| | Tetraploid *mps1AID*-4X* | O755 | O635 * O638 |

### S2 Table: Diploid parent strain list.

***All XX strains***: *ura3-13, trp1-Δ63, his3-Δ1, leu2, met13-d, tyr1-1, lys2-1, can1-R*
***All YY strains***: *ura3-1, trp1-Δ63, his3-Δ1, leu2, met13-c, tyr1-2, lys2-2, cyh2-1*

| | |
|---|---|
| O484 | ***YY strain,*** *MATa/MATa, mps1-R170S::his5/mps1-R170S::his5* |
| O487 | ***YY strain,*** *MAT∝/MAT∝, CEN-plasmid pRS314-mps1-as1* |
| O488 | ***YY strain,*** *MATa/MATa, CEN-plasmid pRS314-mps1-as1* |
| O495 | ***YY strain,*** *MAT∝/MAT∝, mps1-R170S::his5/mps1-R170S::his5, CEN-plasmid pRS314-mps1-as1* |
| O523 | ***YY strain,*** *MAT∝/MAT∝, mps1Δ::KANMX/mps1Δ::KANMX, CEN-plasmid pRS314-mps1-as-1-TRP1* |
| O529 | ***YY strain,*** *MATa/MAT∝, mps1Δ::KANMX/mps1Δ::KANMX, CEN-plasmid pRS314-mps1-as-1-TRP1* |
| O627 | ***XX strain,*** *MATa/MATa CEN3-RS2-HIS3MX6/HphNT1-RS1-CEN3* |
| O629, O630 | ***XX strain,*** *MAT∝/MAT∝, CEN3-RS2-HIS3MX6/HphNT1-RS1-CEN3* |
| O631 | ***XX strain,*** *MATa/ MATa, leu2::[OPL281:LEU2, $P_{GPD1}$-AFB2], leu2::[OPL281:LEU2, $P_{GPD1}$-AFB2], tyr1-1, lys2-1, met13-d, can1-R, CEN3-RS2-HIS3MX6, HphNT1-RS1-CEN3* |
| O633 | ***XX strain,*** *MAT∝/MAT∝, leu2::[OPL281:LEU2, $P_{GPD1}$-AFB2], leu2::[OPL281:LEU2, $P_{GPD1}$-AFB2], tyr1-1, lys2-1, met13-d, can1-R, CEN3-RS2-HIS3MX6, HphNT1-RS1-CEN3* |
| O635 | ***XX strain,*** *MATa/MATa, leu2::[OPL281:LEU2, $P_{GPD1}$-AFB2], leu2::[OPL281:LEU2, $P_{GPD1}$-AFB2], CEN3-RS2-HIS3MX6, HphNT1-RS1-CEN3, MPS1-AID*-9Myc-KAN, MPS1-AID*-9Myc-KAN* |
| O638 | ***XX strain,*** *MAT∝/MAT∝,, CEN3-RS2-HIS3MX6/HphNT1-RS1-CEN3, leu2::[OPL281:LEU2, $P_{GPD1}$-AFB2]/leu2::[OPL281:LEU2, $P_{GPD1}$-AFB2], MPS1-AID*-9Myc-KAN/MPS1-AID*-9Myc-KAN* |
| O639 | ***XX strain,*** *MAT∝/MAT∝, leu2::[OPL281:LEU2, $P_{GPD1}$-AFB2], leu2::[OPL281:LEU2, $P_{GPD1}$-AFB2], CEN3-RS2-HIS3MX6, HphNT1-RS1-CEN3, MPS1-AID*-9Myc-NAT, MPS1-AID*-9Myc-NAT* |
| O753, O754 | ***XX strain,*** *MATa/MATa, CEN3-RS2-HIS3MX6/HphNT1-RS1-CEN3, SPC29-yoEGFP Kan/+* |

### S2 Table: Diploid parent strain list (continued).

| | |
|---|---|
| O749, O750 | ***XX strain,*** *MAT∝/MAT∝, CEN3-RS2-HIS3MX6/HphNT1-RS1-CEN3, SPC29-yoEGFP Kan/+* |
| O849, O850 | ***XX strain,*** *MATa/MATa, CEN3-RS2-HIS3MX6/HphNT1-RS1-CEN3, SPC29-yoEGFP Kan/+, mad2::NATMX6/mad2::NATMX6* |
| O852 | ***XX strain,*** *MAT∝/MAT∝, CEN3-RS2-HIS3MX6/HphNT1-RS1-CEN3, SPC29-yoEGFP Kan/+, mad2::NATMX6/mad2::NATMX6* |
| O573 | ***XX strain,*** *MATa/MATa, mad2::NATMX6/mad2::NATMX6* |
| O575, O576 | ***XX strain,*** *MAT∝/MAT∝, mad2::NATMX6/mad2::NATMX6* |
| O853, O854 | ***XX strain,*** *MATa/MATa, tyr1::[HIS5 pCUP1-AFB2]/tyr1-1, CEN3-RS1-hphNT1/+, SPC29-yoEGFP Kan/+, mps1Δ::KANMX4/+* |
| O855, O856 | ***XX strain,*** *MAT∝/MAT∝, tyr1::[HIS5 pCUP1-AFB2]/tyr1-1, CEN3-RS1-hphNT1/+, SPC29-yoEGFP Kan/+, mps1Δ::KANMX4/+* |

| | |
|---|---|
| O860 | ***XX strain,*** *MAT∝/MAT∝, CEN3-RS2-HIS3MX6/HphNT1-RS1-CEN3, MPS1-AID*-9myc-natNT2/mps1Δ::KANMX4* |
| O857, O858 | ***XX strain,*** *MATa/MATa, CEN3-RS2-HIS3MX6/HphNT1-RS1-CEN3, MPS1-AID*-9myc-natNT2/mps1Δ::KANMX4* |
| O903 | ***XX strain,*** *MATa/MATa, CEN3-RS2-HIS3MX6/HphNT1-RS1-CEN3, lys2::pLL1[PCYC1-GFP-lacI LYS2]/lys2-2, CEN1::pJN2[lacO256 LEU2]/+* |
| O904 | ***XX strain,*** *MAT∝/MAT∝, CEN3-RS2-HIS3MX6/HphNT1-RS1-CEN3, lys2::pLL1[PCYC1-GFP-lacI LYS2]/lys2-2, CEN1::pJN2[lacO256 LEU2]/+* |
| O905, O906 | ***XX strain,*** *MATa/MATa, CEN3-RS2-HIS3MX6/HphNT1-RS1-CEN3, lys2::pLL1[PCYC1-GFP-lacI LYS2]/lys2-2, CEN1::pJN2[lacO256 LEU2]/+, MPS1-AID*-9myc-natNT2/mps1Δ::KANMX4, CEN1::pJN2[lacO256 LEU2]/+* |
| O907, O908 | ***XX strain,*** *MAT∝/MAT∝, CEN3-RS2-HIS3MX6/HphNT1-RS1-CEN3, lys2::pLL1[PCYC1-GFP-lacI LYS2]/lys2-2, CEN1::pJN2[lacO256 LEU2]/+, MPS1-AID*-9myc-natNT2/mps1Δ::KANMX4, CEN1::pJN2[lacO256 LEU2]/+* |
| O924 | ***XX strain,*** *MATa/MATa, leu2::[OPL281:LEU2,* $P_{GPD1}$*-AFB2], leu2::[OPL281:LEU2,* $P_{GPD1}$*-AFB2], CEN3-RS2-HIS3MX6, HphNT1-RS1-CEN3, BUB1-AID*-9myc-KANMX4, BUB1-AID*-9myc-KANMX4, CEN-plasmid pRS314-mps1-as1* |
| O926 | ***XX strain,*** *MAT∝/MAT∝, leu2::[OPL281:LEU2,* $P_{GPD1}$*-AFB2], leu2::[OPL281:LEU2, PGPD-AFB2], CEN3-RS2-HIS3MX6, HphNT1-RS1-CEN3, BUB1-AID*-9myc-KANMX4, BUB1-AID*-9myc-KANMX4, CEN-plasmid pRS314-mps1-as1* |
| O927 | ***XX strain,*** *MATa/MATa, leu2::[OPL281:LEU2,* $P_{GPD1}$*-AFB2], leu2::[OPL281:LEU2,* $P_{GPD1}$*-AFB2], CEN3-RS1-hphNT1, CEN3-RS1-hphNT1, SPC105-AID*-9myc-KANMX4, SPC105-AID*-9myc-KANMX4, his3Δ1::spc105-6A:HIS3, his3Δ1::spc105-6A:HIS3, CEN-plasmid pRS314-mps1-as1* |
| O931 | ***XX strain,*** *MAT∝/MAT∝, leu2::[OPL281:LEU2,* $P_{GPD1}$*-AFB2], leu2::[OPL281:LEU2,* $P_{GPD1}$*-AFB2], CEN3-RS1-hphNT1, CEN3-RS1-hphNT1, SPC105-AID*-9myc-KANMX4, SPC105-AID*-9myc-KANMX4, his3Δ1::spc105-6A:HIS3, his3Δ1::spc105-6A:HIS3, CEN-plasmid pRS314-mps1-as1* |
| O984 | ***XX strain,*** *MATa/MATa, lys2::pLL1[*$P_{CYC1}$*-GFP-lacI LYS2], leu2::[OPL281:LEU2,* $P_{GPD1}$*-AFB2], leu2::[OPL281:LEU2,* $P_{GPD1}$*-AFB2], CEN3-RS2-HIS3MX6, HphNT1-RS1-CEN3, BUB1-AID*-9myc-KANMX4, BUB1-AID*-9myc-KANMX4, +, SPC29-yoEGFP Kan, CEN1::pJN2[lacO256 LEU2]/+, CEN-plasmid pRS314-mps1-as1* |
| O985 | ***XX strain,*** *MATa/MATa, lys2::pLL1[*$P_{CYC1}$*-GFP-lacI LYS2], leu2::[OPL281:LEU2,* $P_{GPD1}$*-AFB2], leu2::[OPL281:LEU2,* $P_{GPD1}$*-AFB2], CEN3-RS2-HIS3MX6, HphNT1-RS1-CEN3, BUB1-AID*-9myc-KANMX4, BUB1-AID*-9myc-KANMX4, +, SPC29-yoEGFP Kan, CEN1::pJN2[lacO256 LEU2]/+, CEN-plasmid pRS314-mps1-as1* |
| O1172 | ***XX strain,*** *MAT∝/ MAT∝, mps1-R170S::his5, mps1-R170S::his5,* OPL571[*CEN3-MPS1-URA3*] |
| O1173 | ***YY strain,*** *MATa/MATa, mps1-R170S::his5/ mps1-R170S::his5,* OPL571[*CEN3-MPS1-URA3*] |
| O1174 | ***YY strain,*** *MAT∝/ MAT∝, mps1-R170S::his5/mps1Δ::KANMX,* OPL571[*CEN3-MPS1-URA3*] |
| O1175 | ***YY strain,*** *MATa/MATa, mps1-R170S::his5/mps1Δ::KANMX,* OPL571[*CEN3-MPS1-URA3*] |
| O1180 | ***XX strain,*** *MATa/MATa, mps1Δ::KANMX/mps1Δ::KANMX,* OPL571[*CEN3-MPS1-URA3*] |
| O1181 | ***YY strain,*** *MATa/MATa, mps1Δ::KANMX/mps1Δ::KANMX,* OPL571[*CEN3-MPS1-URA3*] |
| O1182 | ***YY strain,*** *MAT∝/ MAT, mps1Δ::KANMX/mps1Δ::KANMX,* OPL571[*CEN3-MPS1-URA3*] |

| | |
|---|---|
| O1183 | ***XX strain,*** *MATa/MATa, MPS1/mps1Δ::KANMX,* OPL571[*CEN3-MPS1-URA3*] |
| O1184 | ***XX strain,*** *MATa/MATa, MPS1/mps1Δ::KANMX,* OPL571[*CEN3-MPS1-URA3*] |
| O1185 | ***XX strain,*** *MATa/MATa, mps1Δ::KANMX/mps1Δ::KANMX,* OPL571[*CEN3-MPS1-URA3*] |
| O1186 | ***XX strain,*** *MAT∝/MAT∝, MPS1/mps1Δ::KANMX,* OPL571[*CEN3-MPS1-URA3*] |
| O1187 | ***YY strain,*** *MATa/MATa, MPS1/MPS1,* OPL571[*CEN3-MPS1-URA3*] |
| O1188 | ***YY strain,*** *MAT∝/MAT∝, MPS1/mps1Δ::KANMX,* OPL571[*CEN3-MPS1-URA3*] |
| O1189 | ***YY strain,*** *MAT∝/ MAT∝, mps1Δ::KANMX/mps1Δ::KANMX,* OPL571[*CEN3-MPS1-URA3*] |
| O1190 | ***YY strain,*** *MAT∝/ MAT∝, mps1Δ::KANMX/mps1Δ::KANMX,* OPL571[*CEN3-MPS1-URA3*] |
| O1191 | ***XX strain,*** *MATa/MATa, MPS1/MPS1,* pRS416[*CEN6-URA3*] |
| O1192 | ***YY strain,*** *MAT∝/ MAT∝, MPS1/MPS1,* pRS416[*CEN6-URA3*] |

**Supplemental Table 3: Haploid parent strain list.**

| | |
|---|---|
| X3177 | *MATa, ura3-13, trp1-Δ63, leu2-?, tyr1-1, lys2-1, met13-d, can1-R, SPC29-yoEGFP Kan, CEN3-RS2-HIS3MX6* |
| X3178 | *MAT∝, ura3-13, trp1-Δ63, leu2-?, tyr1-1, lys2-1, met13-d, can1-R, SPC29-yoEGFP Kan, CEN3-RS2-HIS3MX6* |
| X3469 | *MATa, ura3-13, trp1-Δ63, leu2-?, tyr1-1, lys2-1, met13-d, can1-R, SPC29-yoEGFP Kan, CEN3-RS2-HIS3MX6, mad2::NATMX6* |
| X3470 | *MAT∝, ura3-13, trp1-Δ63, leu2-?, tyr1-1, lys2-1, met13-d, can1-R, SPC29-yoEGFP Kan, CEN3-RS2-HIS3MX6, mad2::NATMX6* |
| X3473 | *MATa, ura3-13, trp1-Δ63, leu2-?, tyr1::[HIS5 pCUP1-AFB2], lys2-1, met13-d, can1-R, SPC29-yoEGFP Kan* |
| X3474 | *MAT∝, ura3-13, trp1-Δ63, leu2-?, tyr1::[HIS5 pCUP1-AFB2], lys2-1, met13-d, can1-R, SPC29-yoEGFP Kan* |
| X2833 | *MATa, ura3-13, trp1-Δ63, his3-Δ1, leu2-?, met13-d, tyr1-1, lys2-1, can1-R, HphHNT1-RS1-CEN3, MPS1-AID*-9Myc-NAT* |
| X2834 | *MAT∝, ura3-13, trp1-Δ63, his3-Δ1, leu2-?, met13-d, tyr1-1, lys2-1, can1-R, HphHNT1-RS1-CEN3, MPS1-AID*-9Myc-NAT* |
| X55 | *MATa, ura3-13, trp1-Δ63, his3-Δ1, leu2-?, met13-d, tyr1-1, lys2-1, can1-R* |
| X206 | *MAT∝, ura3-13, trp1-Δ63, leu2, tyr1-1, lys2-1, met13-d, can1-R, his3-Δ1* |
| X258 | *MATa, ura3-13, trp1-Δ63, leu2, tyr1-1, lys2-1, met13-d, can1-R, his3-Δ1, mad2::NATMX6* |
| X259 | *MAT∝, ura3-13, trp1-Δ63, leu2, tyr1-1, lys2-1, met13-d, can1-R, his3-Δ1, mad2::NATMX6* |
| X3493 | *MATa, ura3-13, trp1-Δ63, his3-Δ1, leu2-?, met13-d, tyr1-1, lys2::pLL1[PCYC1-GFP-lacI LYS2], can1-R, CEN1::pJN2[lacO256 LEU2], SPC29-yoEGFP Kan* |
| X3494 | *MATa, ura3-13, trp1-Δ63, his3-Δ1, leu2-?, met13-d, tyr1-1, lys2::pLL1[PCYC1-GFP-lacI LYS2], can1-R, CEN1::pJN2[lacO256 LEU2], SPC29-yoEGFP Kan* |
| X753 | *MATa, can1-R, leu2, lys2-1::pLL1[PCYC1-GFP-lacI LYS2], met13-d, trp1-Δ63, tyr1-1, ura3-13, his3-Δ1, CEN1::pJN2[lacO256 LEU2]* |
| X754 | *MAT∝, can1-R, leu2, lys2-1::pLL1[PCYC1-GFP-lacI LYS2], met13-d, trp1-Δ63, tyr1-1, ura3-13, his3-Δ1, CEN1::pJN2[lacO256 LEU2]* |
| Y2179 | *MAT∝, leu2-?, lys2-2, met13-c, tyr1-2, ura3-1, trp1-Δ63, cyh2-1, his3-Δ1, CEN-plasmid pRS314-mps1-as1-TRP1* |
| Y170 | *MATa, leu2-?, lys2-2, met13-c, tyr1-2, ura3-1, trp1-Δ63, cyh2-1, his3-Δ1* |

| | |
|---|---|
| Y2183 | *MAT∝, leu2-?, lys2-2, met13-c, tyr1-2, ura3-1, trp1-Δ63, cyh2-1, his3-Δ1, CEN-plasmid pRS314-mps1-as1-TRP1, mps1-R170S::his5* |
| Y2181 | *MAT∝, leu2-?, lys2-2, met13-c, tyr1-2, ura3-1, trp1-Δ63, cyh2-1, his3-Δ1, CEN-plasmid pRS314-mps1-as1-TRP1, mps1Δ::KANMX* |
| X3790 | *MATa, ura3-13, trp1-Δ63, leu2-?, tyr1::[HIS5* $P_{CUP1}$*-AFB2], lys2-1, met13-d, can1-R, his3-Δ1, CDC20::[CDC20-AID*-9myc-Kan], BUB1-GFP:natNT2, mps1-R170S::his5* |
| X3799 | *MATa, ura3-13, trp1-Δ63, leu2-?, tyr1::[HIS5* $P_{CUP1}$*-AFB2], lys2-1, met13-d, can1-R, his3-Δ1, CDC20::[CDC20-AID*-9myc-Kan], SPC24-mCherry:Hph, BUB1-GFP:natNT2* |
| X3800 | *MAT∝, ura3-13, trp1-Δ63, leu2-?, tyr1::[HIS5* $P_{CUP1}$*-AFB2], lys2-1, met13-d, can1-R, his3-Δ1, CDC20::[CDC20-AID*-9myc-Kan], SPC24-mCherry:Hph, BUB1-GFP:natNT2* |
| X3804 | *MAT∝, ura3-13, trp1-Δ63, leu2-?, tyr1::[HIS5* $P_{CUP1}$*-AFB2], lys2-1, met13-d, can1-R, his3-Δ1, BUB1-GFP:natNT2, mps1-R170S::his5, SPC24-mCherry:hphNT1* |
| X3805 | *MATa, ura3-13, trp1-Δ63, leu2-?, tyr1::[HIS5* $P_{CUP1}$*-AFB2], lys2-1, met13-d, can1-R, his3-Δ1, CDC20::[CDC20-AID*-9myc-Kan], SPC24-mCherry:hphNT1, BUB1-GFP:natNT2, mps1Δ::KANMX, TRP1::10Xmyc-mps1-as1* |
| X3806 | *MAT∝, ura3-13, trp1-Δ63, leu2-?, tyr1::[HIS5* $P_{CUP1}$*-AFB2], lys2-1, met13-d, can1-R, his3-Δ1, CDC20::[CDC20-AID*-9myc-Kan], SPC24-mCherry:hphNT1, BUB1-GFP:natNT2, mps1Δ::KANMX, TRP1::10Xmyc-mps1-as1* |
| X2608 | *MATa, ura3-13, trp1-Δ63, his3-Δ1, leu2::[OPL281:LEU2,* $P_{GPD1}$*-AFB2], met13-d, tyr1-1, lys2-1, can1-R, hphNT1-RS1-CEN3* |
| X2837 | *MATa, ura3-13, trp1-Δ63, his3-Δ1, leu2::[OPL281:LEU2,* $P_{GPD1}$*-AFB2], met13-d, tyr1-1, lys2-1, can1-R, hphNT1-RS1-CEN3, MPS1-AID*-9Myc-KAN* |
| X3727 | *MATa, ura3-13, trp1-Δ63, his3-Δ1, leu2::OPL281[LEU2* $P_{GPD1}$*-AFB2], met13-d, tyr1-1, lys2-1, can1-R, BUB1-AID*-9myc-KANMX4, CEN3-RS1-hphNT1* |
| O1176 | ***Y strain,*** *MAT∝, ura3-1, tyr1-2, lys2-2, met13-c, trp1-Δ63, leu2-?, his3-Δ1, cyh2-1, leu2, mps1-R170S::his5,* OPL571[*CEN3-MPS1-URA3*] |
| O1177 | ***Y strain,*** *MAT∝, ura3-1, tyr1-2, lys2-2, met13-c, trp1-Δ63, leu2-?, his3-Δ1, cyh2-1, leu2, mps1Δ::KANMX,* OPL571[*CEN3-MPS1-URA3*] |
| O1178 | ***X strain,*** *MATa, ura3-13, tyr1-1, lys2-1, met13-d, can1-R, trp1-Δ63, leu2-?, his3-Δ1, mps1-R170S::his5,* OPL571[*CEN3-MPS1-URA3*] |
| O1179 | ***X strain,*** *MATa, ura3-13, tyr1-1, lys2-1, met13-d, can1-R, trp1-Δ63, leu2-?, his3-Δ1, mps1Δ::KANMX,* OPL571[*CEN3-MPS1-URA3*] |
| O1193 | ***X strain,*** *MATa, ura3-13, trp1-Δ63, his3-Δ1, leu2-?, met13-d, tyr1-1, lys2-1, can1-R, mps1Δ::KANMX,* OPL571[*CEN3-MPS1-URA3*] |
| O1194 | ***Y strain,*** *MAT∝, leu2-?, lys2-2, met13-c, tyr1-2, ura3-1, trp1-Δ63, cyh2-1, his3-Δ1, mps1Δ::KANMX,* OPL571[*CEN3-MPS1-URA3*] |
| O1195 | ***X strain,*** *MATa, ura3-13, trp1-Δ63, leu2-?, tyr1-1, lys2-1, met13-d, can1-R, MPS1,* pRS416[*CEN6-URA3*] |
| O1196 | ***Y strain,*** *MAT∝, ura3-1, trp1-Δ63, leu2-?, tyr1-2, lys2-2, met13-c, cyh2-1, MPS1,* pRS416[*CEN6-URA3*] |
